# PCB-Exposed Gut Microbiota Recapitulate Neurobehavioral Deficits and Reprogram Hippocampal Transcriptomic Signatures

**DOI:** 10.64898/2026.11.05.764721

**Authors:** Youjun P. Suh, Rebecca J. Wilson, Nicole M. Breese, Pamela J. Lein, Hans-Joachim Lehmler, Julia Y. Cui

## Abstract

Polychlorinated biphenyls (PCBs) are persistent environmental pollutants associated with developmental neurotoxicity and autism spectrum disorder (ASD). However, whether PCB-induced alterations in gut microbiota contribute to neurodevelopmental toxicity independently of direct chemical exposure remains unclear. Here, we investigated whether transplantation of PCB-associated gut microbiota produces early neurobehavioral alterations and disrupts brain molecular pathways in offspring. Gut microbiota from conventional dams exposed to the Fox River PCB mixture (FRM; 6.0 mg/kg/day) or vehicle were transplanted into germ-free recipient females without direct PCB administration. At postnatal day 7 (P7), offspring underwent ultrasonic vocalization (USV) testing, whole-brain RNA sequencing (n = 4/sex/group), and untargeted serum metabolomics. Untargeted serum metabolomics were also performed in donor dams. PCB-associated fecal microbiota transplantation (FMT) significantly reduced offspring USVs, reproducing effects observed following direct maternal PCB exposure. Brain transcriptomics in offspring identified substantially more differentially expressed genes in males (9,339) than females (1,082), including 512 shared genes. Concordantly regulated genes revealed downregulation of neurotransmitter signaling and ion channel pathways alongside upregulation of RNA processing pathways. Male-associated responses involved altered protein synthesis and neuronal signaling, whereas female-associated responses involved DNA repair and extracellular matrix remodeling. Despite these molecular differences, behavioral responses showed no significant sex-dependent effect. Maternal metabolomics revealed increased bile acids and decreased indole-3-propionic acid, highlighting potential AhR-NFκB and bile acid-associated signaling mechanisms. Offspring metabolomics further identified alterations in lipid-associated metabolites and metabolic profiles. Collectively, these findings show that transferring PCB-exposed microbiota reproduces early neurobehavioral alterations and induces shared and sex-biased molecular responses, identifying the maternal gut microbiome as a candidate mediator linking PCB exposure to developmental neurotoxicity.

## INTRODUCTION

Autism spectrum disorder (ASD) is a prevalent, lifelong neurodevelopmental condition associated with substantial health, educational, and societal burden worldwide (Baxter et al. 2015; Zeidan et al. 2022). Growing evidence has shown that susceptibility to ASD can be exacerbated by environmental exposure, including polychlorinated biphenyls (PCBs), which are a class of persistent organic pollutants that remain widespread decades after their commercial production was banned in the United States and they were listed for global elimination under the Stockholm Convention (Weber et al. 2018). Their high chemical stability allows PCBs to resist degradation and bioaccumulate in the human body (Liu et al. 2020; Sredlova and Cajthaml 2022). Human exposure therefore continues both through release from legacy reservoirs and through PCBs inadvertently generated as byproducts of contemporary pigment and polymer manufacturing (Anezaki and Nakano 2014; Herkert et al. 2018; Hu and Hornbuckle 2010), whose cumulative mass has been estimated to exceed peak commercial production in the 1970s (Megson et al. 2024). In epidemiological studies, prenatal PCB exposure in humans was associated with lower IQ and reading performance, impaired attention and working memory, and deficits in early childhood cognitive development (Jacobson and Jacobson 1996; 2003; Stewart et al. 2003). Maternal concentrations of selected PCB congeners during pregnancy were further associated with altered odds of ASD and ASD with intellectual disability and with elevated autistic behaviors in early childhood (Lyall et al. 2017), and prenatal PCB exposure was associated with ASD screening risk at 18 months of age in a Japanese birth cohort (Doi et al. 2024). Associations with a clinical ASD diagnosis have nonetheless been inconsistent across cohorts, and maternal PCB concentrations have also been linked to child neurodevelopment indirectly, through placental DNA methylation networks at ASD-associated loci including *CSMD1* and *AUTS2* (Mouat et al. 2023). Moreover, Maternal concentrations of selected PCB congeners during pregnancy have also been associated with altered odds of ASD and ASD with intellectual disability and with elevated autistic behaviors in early childhood (Lyall et al. 2017; Stewart et al. 2003). Consistent with these epidemiological findings, developmental PCB exposure in rodent models alters dendritic growth and experience-dependent plasticity and impaired hippocampal long-term potentiation. Nonhuman-primate studies demonstrated persistent disruption of brain dopaminergic signaling (Gilbert and Crofton 1999; Lein et al. 2007; Seegal et al. 1994; Yang et al. 2009). In addition, developmental exposure to a human-relevant PCB mixture (i.e., the MARBLES PCB mixture) in mice altered ultrasonic vocalizations (USVs), repetitive self-grooming, sociability, and dendritic arborization in sex-, genotype-, dose-, and brain-region-dependent patterns (Keil Stietz et al. 2021; Sethi et al. 2021). Mechanistic studies further demonstrated that PCB exposure dysregulated ryanodine receptor-dependent Ca²⁺ signaling, oxidative stress, and apoptosis (Wayman et al. 2012; Yang and Lein 2010). Each of these mechanisms converges on processes implicated in ASD, including dysregulated intracellular Ca^2+^ signaling, redox imbalance, and altered activity dependent dendritic growth. Together, these findings support PCB exposure as a risk factor for ASD (Keil-Stietz and Lein 2023).

Among various types of environmentally relevant PCB mixtures, the FRM is an environmentally relevant PCB mixture developed to approximate the congener profile found in walleye from the PCB-contaminated Fox River/Green Bay system in northeastern Wisconsin, a historically important source of PCB exposure through consumption of locally caught fish. The FRM consists of 35% Aroclor 1242, 35% Aroclor 1248, 15% Aroclor 1254, and 15% Aroclor 1260 and was formulated specifically to better model human exposure to complex environmental PCB mixtures than individual congeners or a single commercial Aroclor (Kostyniak et al. 2005). Developmental exposure to this mixture is known to produce multiple neurotoxic phenotypes in rodents, including altered response inhibition and operant behavior (Sable et al. 2006), persistent auditory dysfunction (Powers et al. 2006), and disruption of striatal dopamine neurochemistry during development (Dreiem et al. 2010). Consistent with these experimental findings, adolescents from fishing households in the Green Bay region who were at risk of FRM exposure through consumption of fish from the Lower Fox River and other contaminated waters showed associations between higher serum PCB concentrations and poorer cognitive flexibility and visual recognition memory (Sprowles et al. 2022). More recently, we demonstrated that developmental FRM exposure in mice reduced neonatal USVs and impaired social novelty behavior, demonstrating adverse effects on early communication and social behavior relevant to neurodevelopmental disorders (Wilson et al. 2024).

Fecal microbiota transplantation (FMT) is increasingly being used experimentally to test whether gut microbial communities can modify neurological function through the gut microbiota-brain axis. In humans, two case reports described cognitive improvement in patients with Alzheimer disease (AD) after FMT administered for recurrent *Clostridioides difficile* infection (Hazan 2020; Park et al. 2021), while open-label studies and a randomized, double-blind, placebo-controlled trial in children with ASD reported improvements in gastrointestinal symptoms and selected autism-related behavioral measures following microbiota transfer therapies (Kang et al. 2019; Kang et al. 2017; Li et al. 2021; Wan et al. 2024). Preclinical studies in mice similarly show that transfer of healthy microbiota can improve cognition and neuropathology in AD models (Elangovan et al. 2022; Kim et al. 2020) and ameliorate memory and social deficits (Goo et al. 2020). Importantly, however, microbiota transfer can also transmit adverse neurological phenotypes, demonstrating that the neurological consequences of FMT depend on the donor microbial community. Transfer of microbiota from individuals with ASD to germ-free mice induced ASD-relevant behavioral abnormalities and altered brain molecular programs (Sharon et al. 2019), while microbiota from patients with AD impaired cognition and hippocampal neurogenesis in recipient rodents (Grabrucker et al. 2023) and microbiota from AD-model mice similarly impaired memory and neurogenesis in wild-type mice (Kim et al. 2021). Likewise, FMT from chronically stressed mice or from patients with inflammatory disease and depression induced anxiety- and depression-like behaviors accompanied by hippocampal neuroinflammation in recipient mice (Jang et al. 2021; Li et al. 2019; Pu et al. 2022). Together, these bidirectional transplantation studies show that gut microbial communities can be sufficient to either improve or reproduce neurological and neurobehavioral phenotypes, providing a powerful experimental approach for establishing causal links between environmentally or disease-associated gut microbiome alterations and brain function.

Despite growing evidence that developmental PCB exposure and the gut microbiome can independently influence neurobehavioral outcomes, it remains unknown whether PCB-induced gut microbiome alterations mechanistically contribute to developmental neurotoxicity. This gap is particularly important for the environmentally relevant FRM, which has been shown to disrupt early social communication and other neurobehavioral endpoints, yet it is unclear whether these effects require direct chemical exposure or can be transmitted through the altered microbiome itself. If such effects can be transmitted through the altered microbiome rather than requiring direct chemical exposure, neurodevelopmental risk may persist after exposure has ceased and may be only loosely indexed by internal PCB burden, which is the exposure metric on which most epidemiological studies depend. It would also raise the possibility that microbiome directed interventions could reduce risk in exposed populations, although any such inference would first require evidence of transmissibility in humans. The corresponding molecular consequences in the developing brain also remain poorly defined. Therefore, the goal of this study was to determine whether FRM-exposed gut microbiota is sufficient to recapitulate developmental neurobehavioral deficits and induce corresponding brain molecular alterations, including transcriptomic signatures, without direct PCB exposure, thereby providing mechanistic insight into the contribution of the gut microbiome to PCB-associated neurodevelopmental toxicity.

## MATERIALS AND METHODS (note: the experimental design is shown in **Figure 1**)

### Chemicals and reagents

The FRM used to generate the donor microbiota is an environmentally relevant synthetic PCB mixture originally formulated to mimic the congener profile associated with contaminated fish from the Fox River/Green Bay region of Wisconsin. A detailed description of the FRM formulation, chemical authentication, exposure preparation, and developmental dosing paradigm has been reported previously (Kostyniak et al. 2005; Wilson et al. 2024). Other chemicals and reagents, unless otherwise noted, were purchased from Sigma-Aldrich.

**Figure 1.**
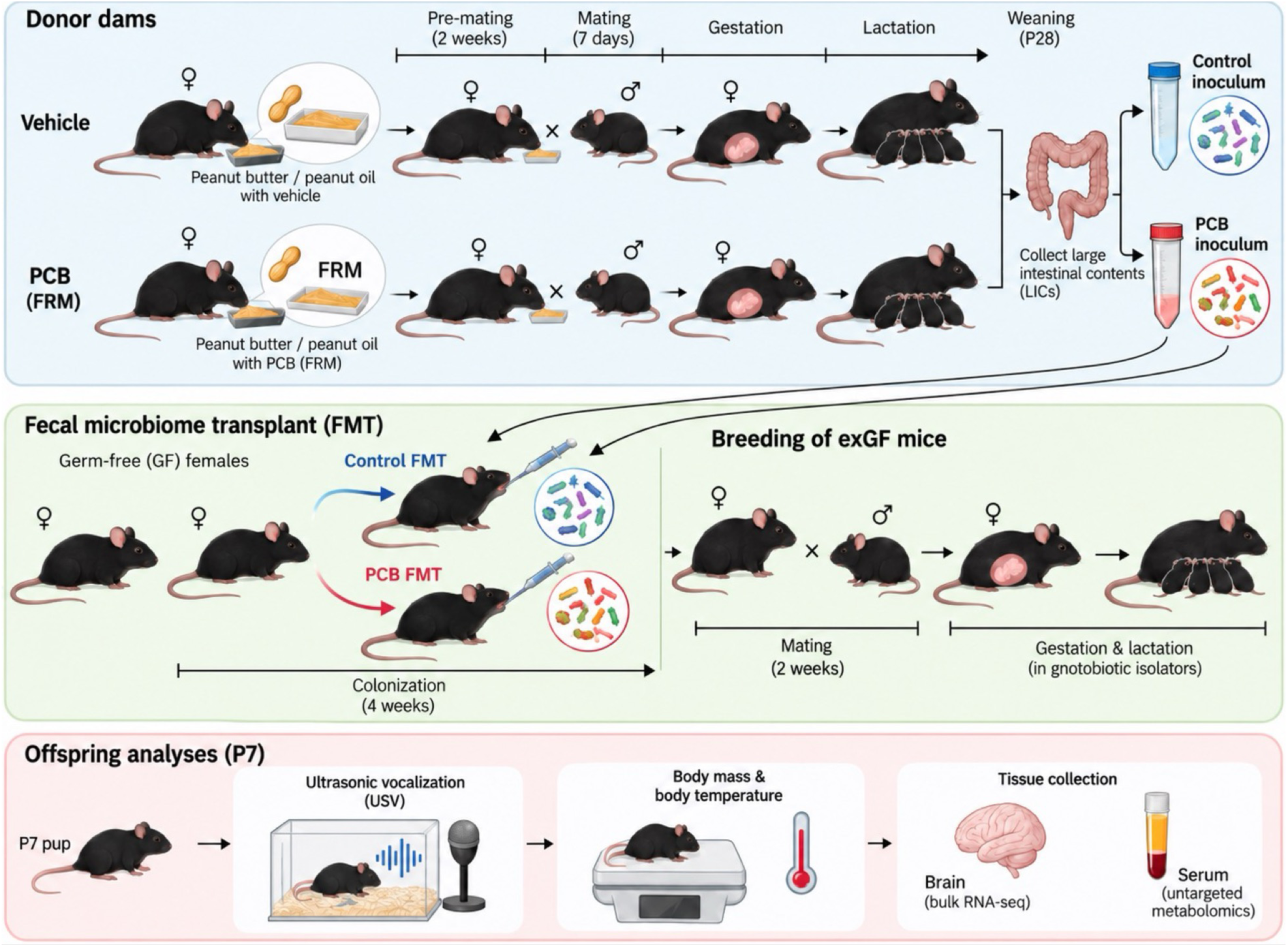
Experimental design of maternal FRM exposure in conventional (CV) C57BL/6 mouse dams, fecal microbiome transplant (FMT), and offspring analyses. Adult conventional (CV) female donor mice were administered vehicle or FRM (6 mg/kg/day) in a peanut butter/peanut oil mixture beginning 2 weeks before mating and continuing through gestation and lactation until weaning at postnatal day 28 (P28). Large intestinal contents (LICs) were collected from donor dams, pooled within treatment group, and used to generate control- and FRM-associated microbiota inocula. Germ-free (GF) female recipient mice received the respective inocula by oral gavage and were maintained under gnotobiotic conditions for 4 weeks to allow microbial colonization. Ex-germ-free (exGF) females were then bred with GF C57BL/6J males and maintained under gnotobiotic conditions throughout gestation and lactation. At P7, offspring underwent ultrasonic vocalization test (USV) followed by body mass and body temperature measurements. For USV test, pups were isolated from the dam, placed into a soundproof chamber, and ultrasonic vocalizations were recorded for 3 minutes. The total numbers of calls during the 3-minute trial were quantified for vehicle and FRM-exposed mice (n=12-19 litters per group, 3-4 pups averaged per litter). Asterisks represent statistically significant differences as compared to the corresponding vehicle groups (*p* < 0.05).

### Animal colony

Conventional (CV) C57BL/6J mice were purchased from The Jackson Laboratory (Bar Harbor, ME, USA) and acclimated to the University of Washington animal facility for 1 week before initiation of experiments. The initial germ-free (GF) C57BL/6 breeding colony was established using mice obtained from the National Gnotobiotic Rodent Resource Center, University of North Carolina (Chapel Hill, NC, USA). Mice were maintained in accordance with Association for Assessment and Accreditation of Laboratory Animal Care International guidelines, and all procedures were approved by the University of Washington Institutional Animal Care and Use Committee (IACUC 4339-01). CV and GF mice were provided the same autoclaved rodent diet (LabDiet #5010; LabDiet, St. Louis, MO, USA), nonacidified autoclaved water, and autoclaved Enrich-N’Pure bedding (The Andersons, Maumee, OH, USA).

### PCB exposure and fecal microbiome transplant

The donor exposure study was conducted at the University of Washington, under an approved Institutional Animal Care and Use Committee protocol (IACUC 4339-01, Cui Laboratory) and in accordance with the NIH *Guide for the Care and Use of Laboratory Animals*. Adult nulliparous CV females (8–10 weeks old) were exposed to vehicle or FRM by oral administration in peanut butter/peanut oil mixture beginning 2 weeks before mating and continuing through gestation and lactation until weaning at postnatal day 28, as described previously (Wilson et al. 2024). The FRM dose was 6.0 mg/kg body weight/day and were adjusted daily based on body weight. This dose was selected on three grounds. Maternal exposure to the FRM at this dose reduced neonatal ultrasonic vocalizations and altered social novelty preference in mice (Wilson et al. 2024); serum PCB and hydroxylated PCB concentrations have been characterized in dams exposed to the same mixture and dosing regimen, providing an internal exposure framework for interpreting biological responses (Li et al. 2024); and maternal dietary exposure at this dose produced brain PCB concentrations comparable to those reported in human brain tissue (Yang et al. 2009). Although the administered dose exceeds typical contemporary environmental intake on a body-weight basis, exposure relevance is more appropriately assessed using internal PCB burdens and congener-specific profiles. The selected regimen therefore provides an established experimental model for investigating microbiome-associated mechanisms of PCB developmental neurotoxicity rather than a direct simulation of typical population-level exposure. After 2 weeks of exposure, an age-matched male was introduced into each female cage for 7 days. After mating, pregnant females were housed individually and remained on their assigned vehicle or FRM exposure throughout gestation and lactation.

For the FMT study, donor large intestinal contents (LICs) were collected from CV dams in the vehicle and 6.0 mg/kg/day FRM groups after weaning of their litters at postnatal day 28. GF recipient mice were not directly exposed to FRM; instead, they were colonized with microbiota derived from these previously exposed donor dams. LICs were flushed from the large intestine with sterile phosphate-buffered saline (PBS) and diluted to approximately 50 mg/mL. Within each treatment group, LICs from all donor dams (n=15 per group) were pooled to generate a single inoculum, yielding two donor preparations, hereafter referred to as vehicle and FRM. Each pooled inoculum was homogenized under anaerobic conditions and passed through a sterile cell strainer to remove large particulate matter. Fresh donor slurry was prepared on the day of transplantation and used immediately without cryopreservation. Each GF recipient female (n=15 per group) received 200 µL of the diluted intestinal-content suspension by oral gavage. Recipients were then maintained under gnotobiotic conditions for 1 month to establish and stabilize the transplanted microbial communities before subsequent breeding and experimental analyses. Recipients colonized with microbiota from FRM-exposed and vehicle-exposed donors are hereafter reffered to as PCB-FMT and vehicle-FMT, respectively, and their offspring as ex-germ-free (exGF) offspring. This FMT procedure has been described previously in our work (Lim et al. 2024).

### Breeding of exGF mice

Following the 4-week colonization period, FMT recipient females were mated with GF C57BL/6J males within their respective gnotobiotic isolators. Mating was limited to a 2-week period to maximize pregnancy success while maintaining a defined gestational window across litters. Pregnant dams remained under gnotobiotic conditions throughout gestation and lactation to maintain the microbial communities established following FMT.

### Ultrasonic vocalization testing

At postnatal day 7 (P7), exGF offspring underwent ultrasonic vocalization (USV) testing to assess early neurobehavioral responses to maternal separation. All pups within each litter were tested because USV responses can vary substantially among littermates. Individual pups were removed from the home cage and placed singly in a container with corn cob bedding inside a sound-attenuating chamber maintained at 22 ± 2°C. USVs were recorded for 3 min using an Avisoft UltraSoundGate microphone and Avisoft Recorder USGH software, version 4.2 (Avisoft Bioacoustics, Glienicke, Germany). Following each recording, pup body mass and body temperature were measured, and pups were returned to the home cage with the dam and littermates. Spectrograms were generated and analyzed using Avisoft SASLab Pro software, version 5.2 (Avisoft Bioacoustics). Recordings were quantified for total call number, call duration, peak frequency, and call type by an investigator blinded to treatment group. For statistical analysis, values from all pups within a litter (3-4 pups per litter) were averaged so that the litter served as the experimental unit (n=15 litters per group).

### Tissue Collection

Following USV testing on P7, exGF offspring and dams were euthanized by CO₂ inhalation, followed by cardiac puncture for blood collection. Seperately, CV donor dams were euthanized, and blood was collected by the same procedure. Serum from these directly exposed donor dams, rather than from exGF recipient dams, was used for the maternal metabolomics reported in Figures 8 and 9. Animals were then perfused with ice-cold sterile PBS to remove residual blood from peripheral and central tissues. Brain tissue was collected immediately after perfusion, snap-frozen in liquid nitrogen, and stored at -80°C until further analysis.

### Total RNA Isolation and Bulk RNA Sequencing

Total RNA was isolated from frozen whole-brain tissue using RNA-Bee reagent (Tel-Test Inc., Friendswood, TX, USA) according to the manufacturer’s protocol. RNA concentration and purity were assessed using a NanoDrop 1000 Spectrophotometer (Thermo Scientific, Waltham, MA, USA) by measuring absorbance at 260 nm. RNA integrity was initially evaluated by formaldehyde-agarose gel electrophoresis with visualization of the 18S and 28S rRNA bands and was further confirmed using an Agilent 2100 Bioanalyzer (Agilent Technologies Inc., Santa Clara, CA, USA). Only RNA samples with an RNA integrity number (RIN) > 8.0 were used for sequencing.

Second-generation bulk RNA sequencing was performed on whole-brain RNA from offspring of dams colonized with either the control or FRM-associated microbiome, with n = 4 per sex per exposure group. Sequencing libraries were prepared using the Illumina TruSeq RNA library preparation kit and sequenced on an Illumina sequencing platform using 150-bp paired-end reads, targeting a sequencing depth of approximately 20 million reads per sample. Raw FASTQ files were aligned to the mouse reference genome GRCm39 using STAR v2.7.11b, with gene annotation from GENCODE release vM35. Gene-level expression data were analyzed separately by sex using limma-voom with sample quality weights. Genes with a mean expression greater than 1 count per million reads (CPM) were retained for analysis.

Differentially expressed genes were defined by an absolute log2 fold change > 1 and a Benjamini-Hochberg false discovery rate (FDR) < 0.05 when comparing offspring of dams colonized with the control versus FRM-associated microbiome.

### Serum Untargeted Metabolomics

During cardiac puncture, whole blood was collected from the heart using a 25-gauge needle and transferred into serum separator tubes. Samples were centrifuged at 2,000 × *g* for 10 min at 4°C, and the resulting serum was collected, snap-frozen in liquid nitrogen, and stored at -80°C until further analysis. Serum samples (dam control, *n =* 12; dam PCB, *n =* 9; male P7 pup control, *n =* 11; male P7 pup PCB, *n =* 9; female P7 pup control, *n =* 10; female P7 pup PCB, *n =* 8) were then subjected to untargeted metabolomics analysis at Novogene (Sacramento, CA, USA). Sample preparation, LC-MS analysis, data processing, and quality control procedures are described in Supplementary Material. A pooled quality control (QC) sample was prepared from all sample extracts to examine the reproducibility of the entire metabolomics process, and one quality control sample was analyzed every 10 test samples. Untargeted metabolomics detected 6472 metabolites, of which 4125 had MS/MS-based structural annotations. Metabolomics data was analyzed via MetaboAnalyst 6.0 online platform (https://www.metaboanalyst.ca/). Initial analyses using principal component analysis and hierarchical clustering were used to evaluate sample-level variation and identify potential outliers. Six serum samples, one from each exposure group, separated from the remaining samples and were excluded from subsequent metabolomics analyses; the sample sizes reported above are those remaining after these exclusions. Data were normalized by quantile, log-transformed, and results were visualized using Partial Least Squares-Discriminant Analysis (PLS-DA), heatmaps, and random forest.

Pathway-level comparison of metabolomic and transcriptomic data. Serum metabolomic and brain transcriptomic datasets were compared at the level of biological pathways rather than by formal statistical integration. For each significantly altered serum metabolite, candidate genes were identified from published pathways in which that metabolite acts as a substrate, product, receptor ligand, or signaling intermediate. Candidate genes were then queried against the sex-specific differential expression results, and those meeting FDR-BH < 0.05 were retained for presentation in Figures 9 and 11. Because candidate genes were selected on the basis of established metabolite-pathway relationships rather than by unbiased statistical association, no additional multiple-testing correspondences are descriptive and hypothesis-generating rather than evidence of directional or causal relationships.

## RESULTS

### PCB-associated gut microbiota reproduced early-life neurobehavioral alterations observed following direct maternal PCB exposure

Early postnatal communication between mouse pups and the dam utilizes ultrasonic vocalizations (USVs), which are emitted by pups when they are separated from the nest (Simola and Granon 2019). These high-frequency calls stimulate maternal behaviors such as searching, approach, and retrieval.(Hahn and Lavooy 2005) Because the frequency and pattern of isolation-induced USVs are sensitive to disruptions in neurobehavioral development, they are widely used as an endpoint for evaluating early communicative behavior in rodent models.(Premoli et al. 2023) Male and female pups did not exhibit significantly different responses as measured by a two-way ANOVA with sex and exposure group as variables (**Supplemental Figure 1A**). Therefore, male and female data were pooled, and the average value of the entire litter was used as the statistical unit. Pups from the PCB-FMT group emitted significantly fewer ultrasonic vocalizations after maternal separation than pups from the vehicle-FMT group (**Figure 2**), indicating impaired early social communication. Notably, the reduction in ultrasonic vocalizations was like that previously observed after direct maternal exposure to 6.0 mg/kg/d of the FRM (**Figure 2**). Together, these findings suggest that transferring a PCB-associated gut microbiome is sufficient to reproduce aspects of the early neurobehavioral effects observed after direct PCB exposure.

**Figure 2.**
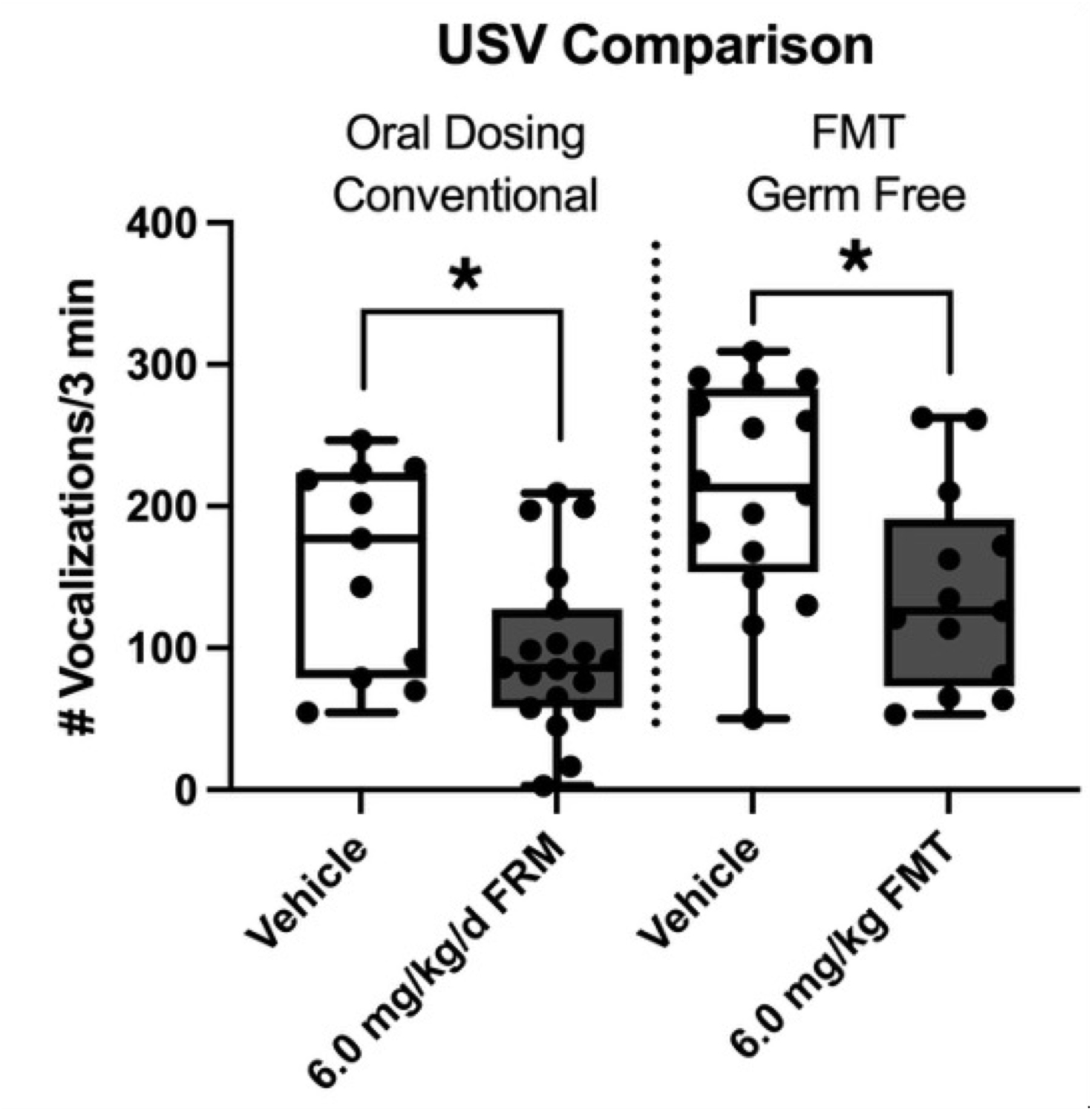
Maternal transfer of FRM-associated gut microbiota recapitulates PCB-induced deficits in early social communication in P7 offspring. Left: CV C57BL/6 dams were exposed to vehicle or the Fox River PCB Mixture (FRM; 6 mg/kg/day) in a peanut butter/peanut oil mixture, and ultrasonic vocalizations (USVs) were assessed in their offspring at P7. Right: GF recipient females were colonized by fecal microbiome transplantation (FMT) with microbiota obtained from vehicle- or FRM-exposed CV mice, as described in Figure 1. The recipient females were not directly exposed to FRM. At P7, exGF offspring born to dams colonized with FRM-associated microbiota exhibited a similar reduction in USV calls compared with offspring receiving control microbiota, demonstrating that the FRM-associated gut microbiota was sufficient to reproduce the early social communication phenotype associated with maternal FRM exposure. For USV test, individual P7 pups were separated from the dam and placed in a sound-attenuating chamber, and vocalizations were recorded for 3 min. Total call number was quantified, with 3–4 pups averaged per litter and the litter used as the experimental unit (n = 12–19 litters per group). Asterisks indicate significant differences relative to the corresponding control group (*p* < 0.05).

To determine whether variation in ultrasonic vocalizations was associated with physiological factors, we examined the relationships between USV counts and body weight or body temperature in postnatal day 7 (P7) offspring. As expected, body weight differed significantly between male and female offspring in both the vehicle-FMT (P < 0.0001) and PCB-FMT (P = 0.0007) groups (**Supplemental Figure 1B**), whereas body temperature was generally comparable across groups (**Supplemental Figure 1C**). Linear regression analyses revealed no significant association between USV counts and body weight (R² = 0.0398, p = 0.2904; **Supplemental Figure 1D**) or body temperature (R² = 0.0007, P = 0.8888; **Supplemental Figure 1E**). Body weight and temperature explained only 3.98% and 0.07% of the variation in USV counts, respectively. These findings suggest that the reduction in USVs observed following PCB-FMT is unlikely to be explained solely by differences in body weight or temperature, supporting an association with altered early-life vocal communication rather than these physiological parameters.

### PCB-FMT induces pronounced sex-dependent brain transcriptional alterations involving autism-associated genes

To determine the brain transcriptomic response to PCB-associated gut microbiota FMT, including neurodevelopment-related gene expression, we performed whole-brain bulk RNA sequencing in postnatal day 7 (P7) male and female offspring following maternal FMT with PCB-exposed or vehicle-exposed donor mice. Differential expression analysis (FDR-BH < 0.05) identified substantially more differentially expressed genes (DEGs) in males than females (**Figure 3A**). Specifically, 9,339 DEGs were identified in male offspring, including 5,347 upregulated and 3,992 downregulated genes. In contrast, female offspring exhibited 1,082 DEGs, comprising 956 upregulated and 126 downregulated genes. Thus, male offspring displayed approximately 8.6-fold more DEGs than females, suggesting a broader transcriptional response to PCB-FMT.

**Figure 3.**
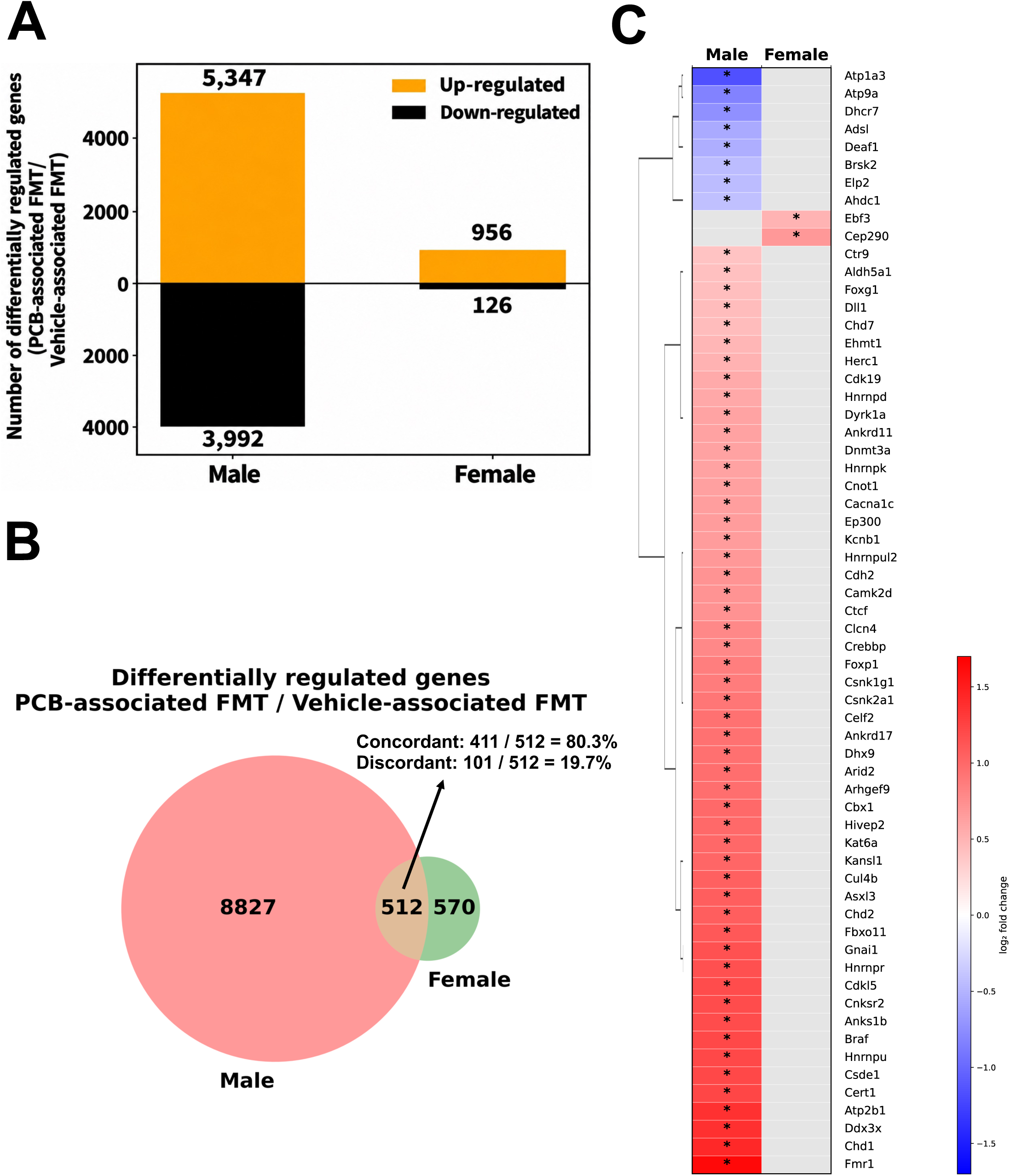
PCB-associated FMT induces sex-dependent brain transcriptional alterations involving ASD associated genes in P7 offspring. Whole-brain RNA sequencing was performed using n = 4 biological replicates per sex per treatment group (male vehicle-FMT, n = 4; male PCB-FMT, n = 4; female vehicle-FMT, n = 4; female PCB-FMT, n = 4; total n = 16). Differential gene expression was analyzed separately by sex using limma-voom with sample quality weights. Statistical significance was defined as a Benjamini–Hochberg-adjusted *P* < 0.05 (false discovery rate [FDR-BH] < 0.05). Genes significant in only one sex were classified as sex-specific differentially regulated genes (DEGs) for visualization. (**A**) Numbers of DEGs in male and female P7 offspring brains following PCB-associated FMT compared with vehicle-associated FMT. Orange and black bars indicate upregulated and downregulated genes, respectively. (**B**) Venn diagram showing the overlap of DEGs between male and female offspring. (**C**) Hierarchically clustered heatmap showing 62 selected ASD-associated DEGs identified by cross-referencing significantly regulated offspring brain genes with the Simons Foundation Autism Research Initiative (SFARI) Gene database (https://gene.sfari.org/; 2026 Q2 reference subset). Colors indicate log2 fold changes (red, upregulation; blue, downregulation), and asterisks (*) indicate statistically significant differential expression. Gray cells represent genes not identified as significant in the corresponding sex in the analyzed DEG lists.

Comparison of the male and female DEG profiles identified 512 shared genes, whereas 8,827 and 570 DEGs were uniquely identified in males and females, respectively (**Figure 3B**). Among the shared DEGs, 411 genes (80.3%) exhibited concordant regulation in both sexes, whereas 101 genes (19.7%) were regulated in opposite directions. These findings indicate substantial sex differences in the magnitude and composition of the transcriptional response, along with considerable directional agreement among shared DEGs.

To investigate the potential relevance of these transcriptional alterations to autism spectrum disorder (ASD), we cross-referenced the offspring brain DEGs with ASD-associated genes curated in the Simons Foundation Autism Research Initiative (SFARI) Gene database (https://gene.sfari.org/). This comparison identified 62 ASD-associated DEGs: 60 significant in males and 2 significant in females (**Figure 3C**). Hierarchical clustering of their log2 fold changes revealed a predominantly male-associated transcriptional response, with both upregulated and downregulated genes. Notably, PCB-FMT induced significant upregulation of several neurodevelopmental genes in male offspring, including *Fmr1* (fragile X messenger ribonucleoprotein 1), *Chd1* (chromodomain helicase DNA-binding protein 1), *Chd2* (chromodomain helicase DNA-binding protein 2), *Cdkl5* (cyclin-dependent kinase-like 5), *Foxg1* (forkhead box G1), *Foxp1* (forkhead box P1), *Dyrk1a* (dual-specificity tyrosine-phosphorylation-regulated kinase 1A), *Dnmt3a* (DNA methyltransferase 3 alpha), and *Ankrd11* (ankyrin repeat domain-containing protein 11). These genes are annotated in SFARI Gene as functioning in chromatin remodeling, epigenetic regulation, neuronal differentiation, synaptic plasticity, and brain development. In contrast, *Atp1a3* (ATPase Na⁺/K⁺ transporting subunit alpha 3), *Atp9a* (ATPase phospholipid transporting 9A), *Dhcr7* (7-dehydrocholesterol reductase), and *Adsl* (adenylosuccinate lyase) were downregulated in males, suggesting potential alterations in neuronal excitability, membrane lipid trafficking, cholesterol biosynthesis, and purine metabolism.

In female offspring, only two genes from the SFARI-curated set were significantly differentially expressed: *Ebf3* (early B-cell factor 3), and *Cep290* (centrosomal protein 290) (Figure3C). *Ebf3* is annotated as a transcription factor functioning in neuronal differentiation and development, and *Cep290* as a centrosomal protein required for primary cilium assembly. Collectively, these findings suggest that PCB-associated gut microbiota perturb transcriptional programs linked to neurodevelopment and ASD susceptibility, with more extensive alterations observed in the male offspring.

### PCB-FMT alters neuronal signaling and RNA processing pathways in both male and female offspring

To identify molecular pathways commonly affected by PCB-FMT in both sexes, we performed Enrichr Reactome pathway enrichment analyses of genes concordantly downregulated or upregulated in P7 offspring brains compared with vehicle-associated FMT controls.

Concordantly downregulated genes by PCB-FMT in brains of both male and female pups were enriched in pathways associated with neuronal neurotransmission and cellular stress responses (**Figure 4A**). Notably, multiple pathways involved in nicotinic acetylcholine receptor signaling were represented, including highly sodium-permeable postsynaptic nicotinic acetylcholine receptors (P = 0.0143), highly calcium-permeable nicotinic acetylcholine receptors (P = 0.0183), presynaptic nicotinic acetylcholine receptors (P = 0.0243), and acetylcholine binding and downstream signaling events (P = 0.0283). Additional enriched pathways included adrenergic receptors (P = 0.0183), tandem pore domain potassium channels (P = 0.0243), and nuclear factor erythroid 2-related factor 2 (NFE2L2)-regulated multidrug resistance-associated enzymes (P = 0.0163). These findings suggest that PCB-FMT is associated with coordinated downregulation of genes involved in cholinergic and adrenergic neurotransmission, ion channel function, and cellular stress-response pathways in both sexes. Several of these pathways converge on Ca^2+^ dependent neuronal signaling, including the highly calcium-permeable nicotinic acetylcholine receptor pathway identified here. Direct PCB exposure has been reported to dysregulate ryanodine receptor-dependent Ca2+ release from intracellular stores (Wayman et al. 2012; Yang and Lein 2010), whereas the present findings implicate transcriptional downregulation of receptor-and channel-mediated Ca^2+^ entry; whether these converge functionally remains to be determined, but both identify Ca^2+^ dependent signaling as a point of vulnerability in the developing brain.

**Figure 4.**
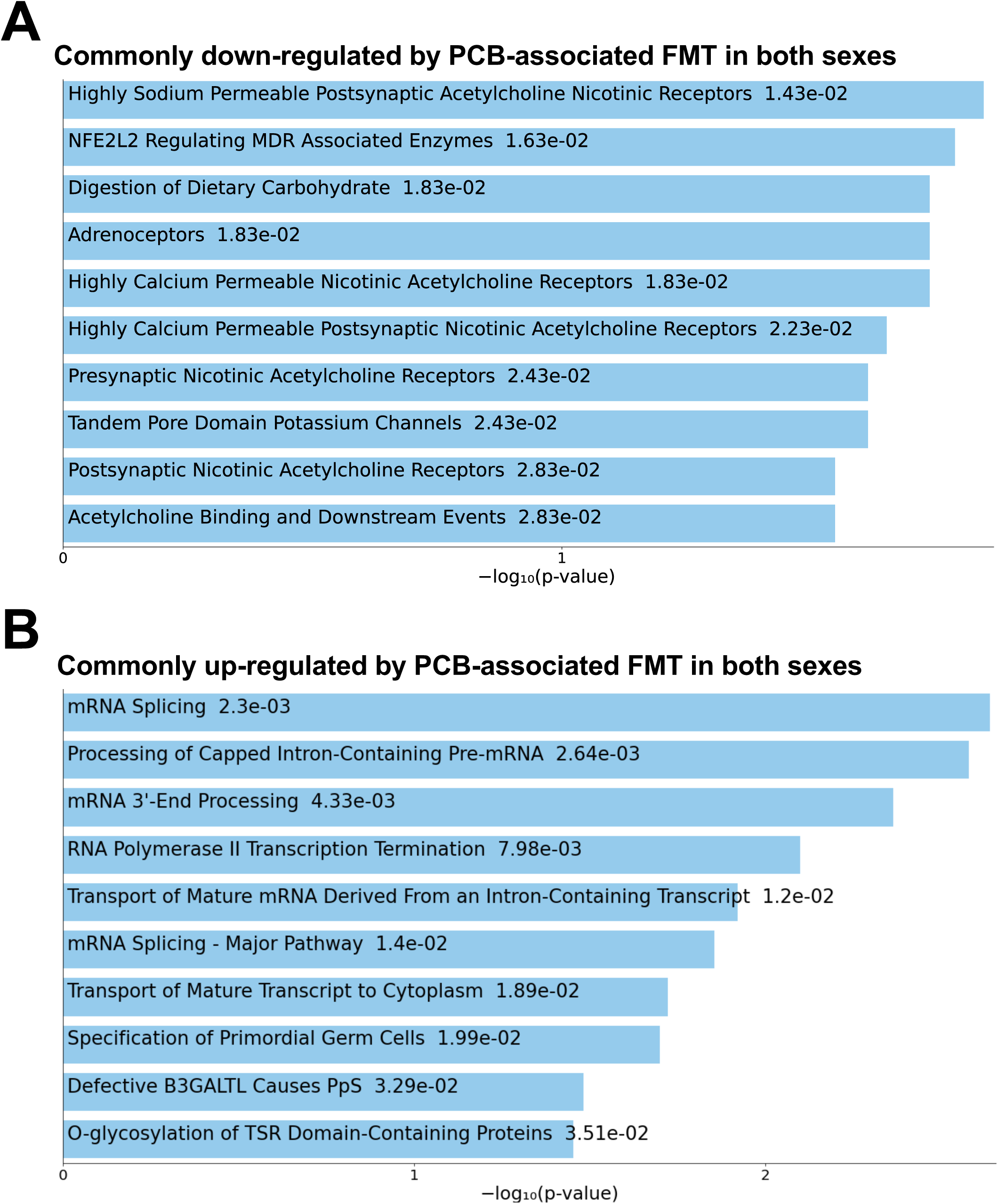
PCB-associated FMT alters neuronal signaling and RNA processing pathways in both male and female P7 offspring brains. **(A)** Reactome pathway enrichment analysis of 41 genes concordantly downregulated in male and female P7 offspring brains following PCB-associated FMT compared with vehicle-associated FMT. Whole-brain RNA sequencing was performed using n = 4 biological replicates per sex per FMT group (total n = 16). Differential gene expression was analyzed separately by sex using limma-voom with sample quality weights and FDR-BH < 0.05, as described in Figure 3. Significantly downregulated genes shared between sexes were analyzed using Enrichr with the Reactome pathway database. **(B)** Reactome pathway enrichment analysis of 370 genes concordantly upregulated in both sexes, identified using the same RNA-seq sample sizes and statistical criteria described in (**A**). Enrichment analysis was performed using Enrichr with the Reactome pathway database. For both panels, pathways were ranked by enrichment significance, and values displayed alongside pathway names represent Enrichr enrichment *P* values.

In contrast, concordantly upregulated genes by PCB-FMT in brains of both male and female pups were predominantly enriched in pathways involved in mRNA processing and post-transcriptional regulation (**Figure 4B**). The most significantly enriched pathways included mRNA splicing (P = 0.0023), processing of capped intron-containing precursor mRNA (P = 0.00264), mRNA 3′-end processing (P = 0.00433), and RNA polymerase II transcription termination (P = 0.00798). Additional pathways involved major-pathway mRNA splicing (P = 0.014), transport of mature mRNA derived from intron-containing transcripts (P = 0.012), and transport of mature transcripts to the cytoplasm (P = 0.0189). These results indicate that PCB-FMT increased expression of genes involved in RNA maturation, processing, and transport in developing offspring brains.

### Discordantly regulated genes reveal sex-opposing alterations in inflammatory signaling, nuclear receptor regulation, extracellular matrix organization, and neurite development

Although PCB-FMT induced substantial transcriptional alterations that are concordant in both male and female P7 offspring brains, a small fraction of shared differentially expressed genes (DEGs) exhibited opposite directions of regulation between sexes (101 genes, 19.7%). Of these 101 discordant genes, 29 were upregulated in males but downregulated in females, while 72 were downregulated in males but upregulated in females.

As shown in **Figure 5A**, Reactome analysis of the 29 genes upregulated in males but downregulated in females revealed enrichment of pathways involved in intracellular receptor regulation, inflammatory signaling, and lipid mediator metabolism. The most prominently enriched pathways included SUMOylation of intracellular receptors (P = 8.61 × 10⁻□), interleukin-10 (IL-10) signaling (P = 2.02 × 10⁻³), nuclear receptor transcription (P = 2.67 × 10⁻³), O-linked glycosylation of mucins (P = 3.99 × 10⁻³), and G protein-coupled receptor (GPCR) ligand binding (P = 4.39 × 10⁻³). Additional enriched pathways included synthesis of 15-eicosatetraenoic acid derivatives, biosynthesis of eicosapentaenoic acid (EPA)-derived specialized pro-resolving mediators (SPMs), and regulation of estrogen receptor- and WNT-associated transcriptional pathways. These findings suggest sex-opposing transcriptional responses involving receptor-mediated signaling, immune regulation, and lipid-derived inflammatory mediators. Notably, the enrichment of IL-10 signaling, which is positively linked to autism in humans (Saghazadeh et al. 2019), as well as the enrichment of SPM biosynthesis pathways, which is positively linked to inflammation resolution in ASD (Basil and Levy 2016; Sumra et al. 2025; Valente et al. 2022), raises the possibility of differential regulation of anti-inflammatory or inflammation-resolving processes between male and female offspring.

**Figure 5.**
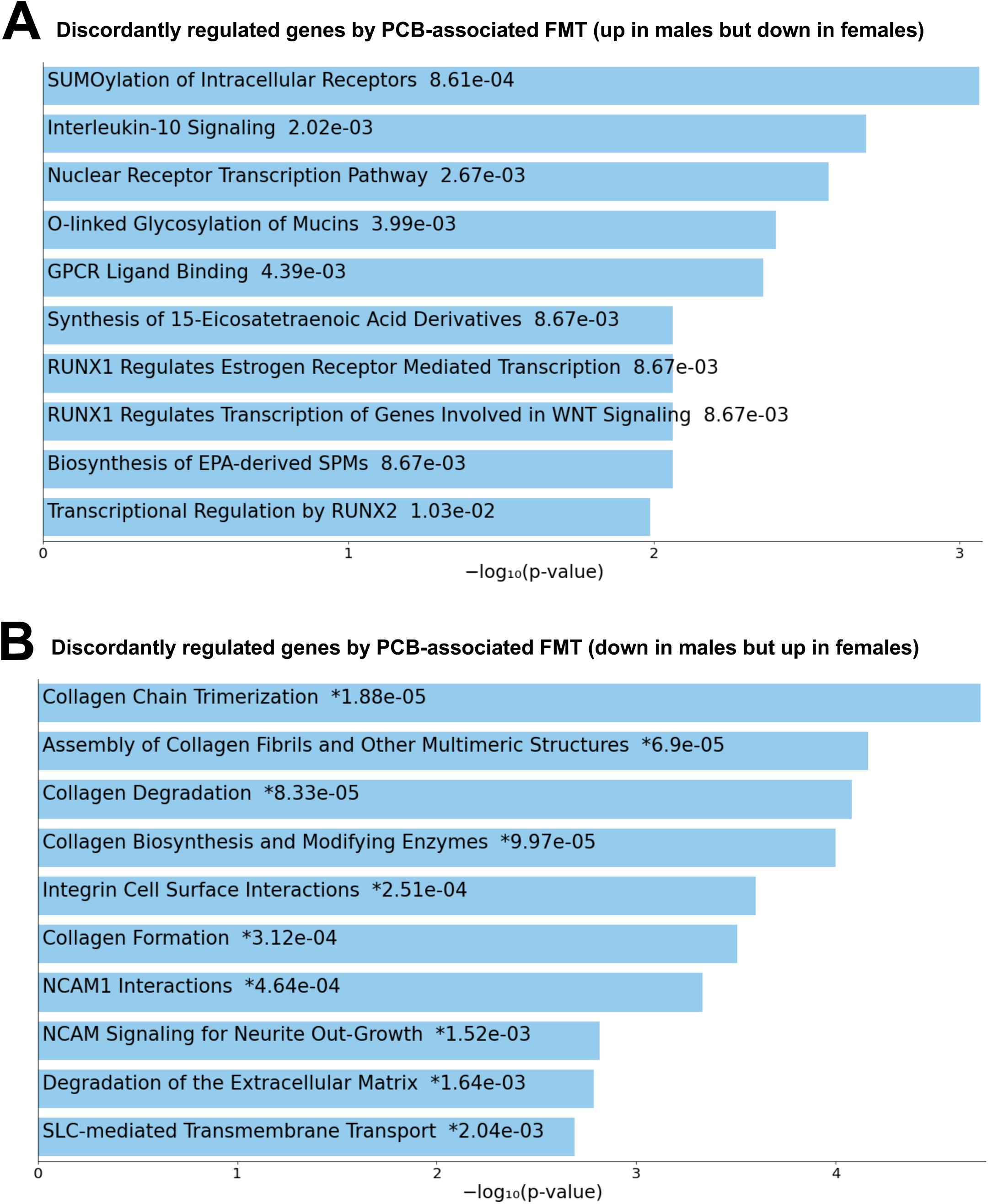
PCB-associated FMT induces sex-opposing transcriptional alterations involving inflammatory signaling, nuclear receptor regulation, extracellular matrix organization, and neurite development in P7 offspring brains. (A) Reactome pathway enrichment analysis of 29 genes significantly upregulated in male but downregulated in female P7 offspring brains following PCB-associated FMT compared with vehicle-associated FMT. Whole-brain RNA sequencing was performed using n = 4 biological replicates per sex per FMT group (total n = 16). Differential gene expression was analyzed separately by sex using limma-voom with sample quality weights and FDR-BH < 0.05, as described Figure 3. The 29 discordantly regulated genes were subjected to pathway enrichment analysis using Enrichr with the Reactome pathway database. (B) Reactome pathway enrichment analysis of 72 genes significantly downregulated in males but upregulated in females, identified using the same RNA-seq sample sizes and statistical criteria described in (A). Enrichment analysis was performed using Enrichr with the Reactome pathway database. For both panels, the top 10 pathways were ranked by enrichment significance, and values displayed alongside pathway names represent Enrichr enrichment *P* values. Asterisks (*) indicate pathways marked as significant after multiple-testing correction in the enrichment output, where applicable.

Conversely, as shown in **Figure 5B**, the 72 genes downregulated in males but upregulated in females were significantly enriched in pathways involved in extracellular matrix (ECM) organization, collagen metabolism, cell adhesion, and neuronal development. These included collagen chain trimerization (P = 1.88 × 10⁻□), assembly of collagen fibrils and other multimeric structures (P = 6.90 × 10⁻□), collagen degradation (P = 8.33 × 10⁻□), collagen biosynthesis and modifying enzymes (P = 9.97 × 10⁻□), and integrin cell surface interactions (P = 2.51 × 10⁻□). Importantly, neural cell adhesion molecule 1 (NCAM1) interactions (P = 4.64 × 10⁻□) and NCAM signaling for neurite outgrowth (P = 1.52 × 10⁻³) were also enriched. These pathways are important for developmental neurotoxicity because ECM organization, cell adhesion, and NCAM-mediated signaling contribute to neuronal migration, neurite extension, synapse formation, and neural circuit maturation (Duncan et al. 2021). The opposite regulation of these genes in males and females suggests distinct transcriptional responses that may affect structural and developmental processes in the early postnatal brain.

### Male-specific transcriptional alterations reveal suppression of protein synthesis and enrichment of neuronal signaling pathways following PCB-FMT

To further characterize the sex-specificity in the brain transcriptomic response to PCB-FMT, Reactome pathway enrichment analysis was performed on 8,827 DEGs identified only in male but not female P7 offspring brains, including 3,879 downregulated (**Figure 6A**) and 4,948 upregulated genes (**Figure 6B**).

**Figure 6.**
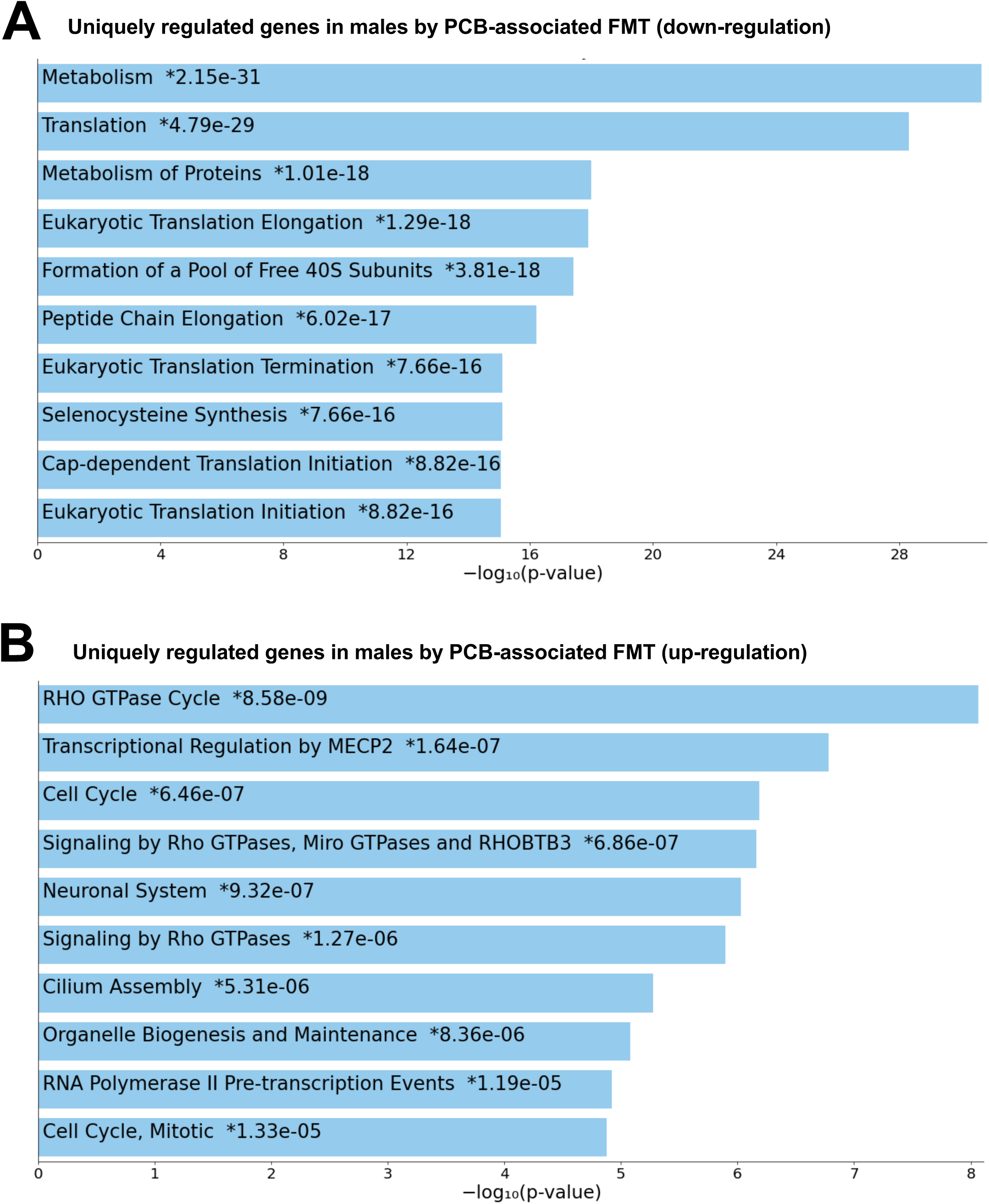
PCB-associated FMT induces male-specific brain transcriptional alterations involving protein synthesis, neuronal signaling, and cytoskeletal regulation in P7 offspring. (A) Reactome pathway enrichment analysis of 3,879 genes significantly downregulated in male but not female P7 offspring brains following PCB-associated FMT compared with vehicle-associated FMT. Whole-brain RNA sequencing was performed using n = 4 biological replicates per sex per FMT group (total n = 16). Differential gene expression was analyzed separately by sex using limma-voom with sample quality weights and FDR-BH < 0.05, as described Figure 3. Male-specific downregulated genes were subjected to pathway enrichment analysis using Enrichr with the Reactome pathway database. (B) Reactome pathway enrichment analysis of 4,948 genes significantly upregulated in male but not female offspring brains, identified using the same RNA-seq sample sizes and statistical criteria described in (**A**). Enrichment analysis was performed using Enrichr with the Reactome pathway database. For both panels, the top 10 enriched pathways were ranked by enrichment significance, and values displayed alongside pathway names represent Enrichr enrichment p values. Asterisks (*) indicate statistically significant pathways. Male-specific genes refer to DEGs reaching statistical significance only in males.

As shown in **Figure 6A**, male-specific downregulated genes were significantly enriched in pathways involved in metabolism (P = 2.15 × 10⁻³¹), translation (P = 4.79 × 10⁻²□), protein metabolism (P = 1.01 × 10⁻¹□), and multiple processes associated with ribosomal function and protein synthesis. These findings suggest broad transcriptional suppression of pathways essential for neuronal growth, differentiation, and synaptic development. To note, dysregulation of these pathways has been implicated in ASD and other neurodevelopmental disorders (Joo and Benavides 2021; Santini and Klann 2014).

In contrast, male-specific upregulated genes (**Figure 6B**) were enriched in the RHO GTPase cycle (P = 8.58 × 10⁻□), transcriptional regulation by methyl-CpG-binding protein 2 (MECP2; P = 1.64 × 10⁻□), cell cycle (P = 6.46 × 10⁻□), neuronal system (P = 9.32 × 10⁻□), and cilium assembly (P = 5.31 × 10⁻□). These pathways regulate cytoskeletal dynamics, neuronal differentiation, synaptic organization, and neural circuit maturation, processes whose dysregulation has been implicated in neurodevelopmental disorders, including ASD (Falace et al. 2026; Guo et al. 2020).

### Female-specific transcriptional alterations reveal downregulation of neurotransmitter signaling and upregulation of DNA repair and extracellular matrix pathways following PCB-FMT

To characterize female-specific transcriptional responses to PCB-associated fecal microbiota transplantation (FMT), Reactome pathway enrichment analysis was performed on 570 differentially expressed genes (DEGs) identified in female but not male P7 offspring brains, including 56 downregulated (**Figure 7A**) and 514 upregulated genes (**Figure 7B**).

**Figure 7.**
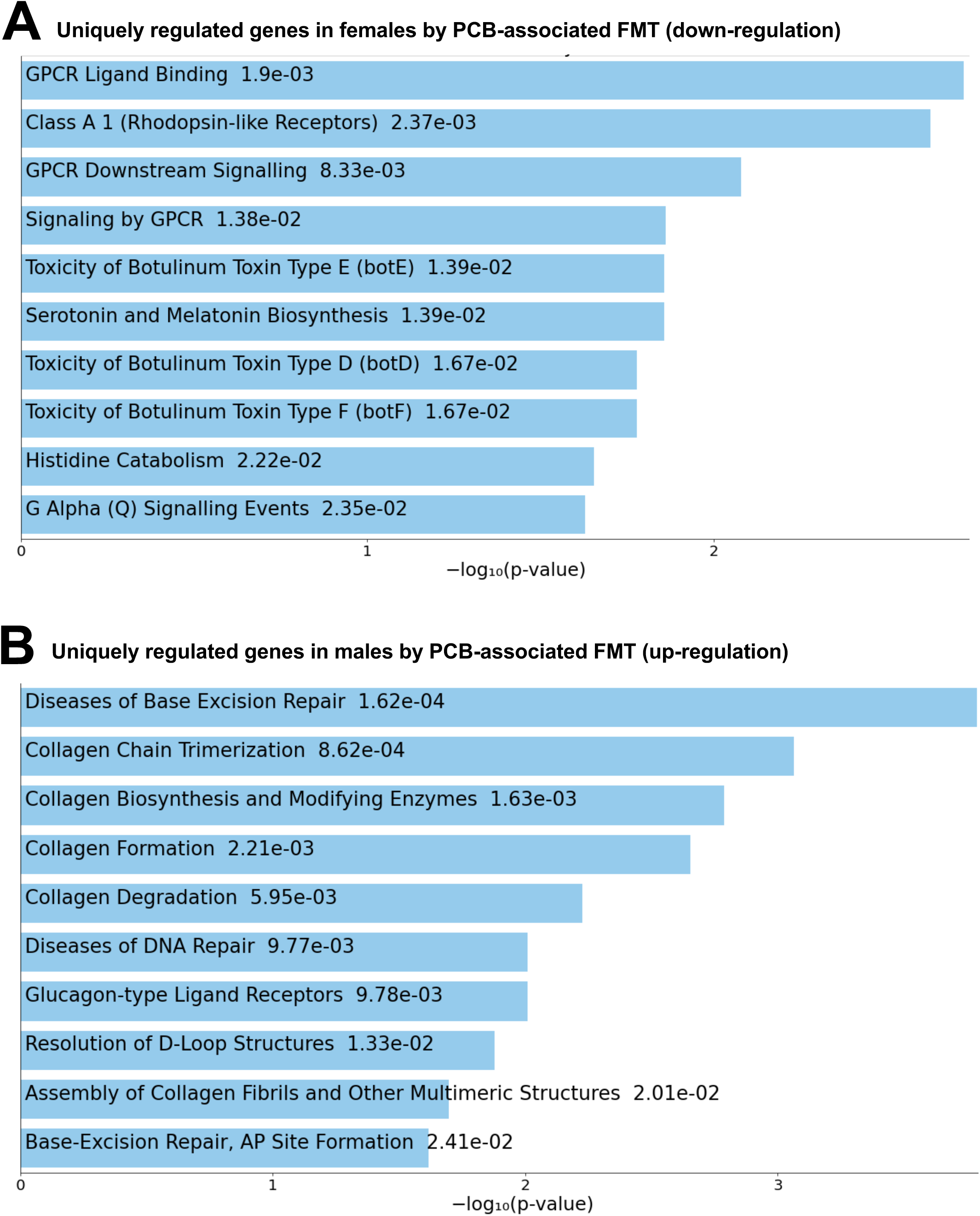
PCB-associated FMT induces female-specific brain transcriptional alterations involving neurotransmitter signaling, DNA repair-associated pathways, and extracellular matrix remodeling in P7 offspring. (A) Reactome pathway enrichment analysis of 56 genes significantly downregulated in female but not male P7 offspring brains following PCB-associated FMT compared with vehicle-associated FMT. Whole-brain RNA sequencing was performed using n = 4 biological replicates per sex per FMT group (total n = 16). Differential gene expression was analyzed separately by sex using limma-voom with sample quality weights and FDR-BH < 0.05, as described in Figure 3. Female-specific downregulated genes were subjected to pathway enrichment analysis using Enrichr with the Reactome pathway database. **(B)** Reactome pathway enrichment analysis of 514 genes significantly upregulated in female but not male offspring brains, identified using the same RNA-seq sample sizes and statistical criteria described in (**A**). Enrichment analysis was performed using Enrichr with the Reactome pathway database. For both panels, the top 10 enriched pathways are shown, ranked by enrichment significance. Values displayed alongside pathway names represent Enrichr enrichment P values. Female-specific genes refer to DEGs reaching statistical significance only in females. Disclaimer: Generative AI was used only for figure graphics, not for scientific ideas, experimental design, or data.

As shown in **Figure 7A**, female-specific downregulated genes were enriched in GPCR ligand binding (P = 1.90 × 10⁻³), class A/1 rhodopsin-like receptors (P = 2.37 × 10⁻³), GPCR downstream signaling (P = 8.33 × 10⁻³), and serotonin and melatonin biosynthesis (P = 1.39 × 10⁻²). These findings suggest altered receptor-mediated neurotransmission and neurochemical signaling, processes essential for neuronal communication, synaptic development, and brain maturation. To note, the dysregulation of these pathways has been implicated in ASD (Annamneedi et al. 2025).

In contrast, female-specific upregulated genes (**Figure 7B**) were enriched in pathways associated with base excision repair (P = 1.62 × 10⁻□), collagen chain trimerization (P = 8.62 × 10⁻□), collagen biosynthesis and modification (P = 1.63 × 10⁻³), collagen formation (P = 2.21 × 10⁻³), and collagen degradation (P = 5.95 × 10⁻³). These pathways are involved in DNA damage repair and ECM organization, contributing to genomic stability, neuronal migration, neurite extension, and synaptic maturation. Their upregulation may reflect altered tissue remodeling and potentially adaptive DNA repair-associated transcriptional responses. Notably, disruptions in DNA repair and ECM organization have been implicated in neurodevelopmental disorders, including autism spectrum disorder (Kelvington et al. 2026; Wen et al. 2018).

### Maternal PCB exposure alters circulating bile acids, microbiome-derived metabolites, and oxidative stress-associated pathways

To explore potential metabolic mechanisms underlying the neurobehavioral and transcriptomic alterations observed following PCB-FMT, we next examined serum metabolomic profiles of PCB-exposed donor dams. Because gut microbiota play important roles in xenobiotic and endogenous metabolism (Klaassen and Cui 2015; Koppel et al. 2017), PCB-induced alterations in maternal circulating metabolites may provide insights into microbiome-associated metabolic pathways potentially contributing to offspring neurodevelopmental effects.

As shown in **Figure 8A**, untargeted serum metabolomics was performed in CV donor dams exposed to vehicle or the FRM (6.0 mg/kg/day), the same animals whose large intestinal contents were used to generate the FMT inocula. FRM exposure significantly altered 92 circulating metabolites, including 68 increased and 24 decreased metabolites. Pathway enrichment analysis highlighted porphyrin metabolism, with 9 of 31 pathway-associated metabolites represented (**Figure 8B**), suggesting potential alterations in heme-related metabolic processes.

Further examination of individual metabolites revealed significant increases in several bile acid species, including β-muricholic acid (P = 0.0476), hyocholic acid (P = 0.0395), ursocholic acid (P = 0.0200), ursocholytaurine (P = 0.0457), and glycochenodeoxycholic acid 7-sulfate (P = 0.0365) (**Figure 8C**). Given the established roles of bile acids in receptor-mediated signaling, metabolic homeostasis, and neuroimmune regulation (Joyce and O’Malley 2022; Monteiro-Cardoso et al. 2021), these changes may be relevant to offspring neurodevelopment. Although altered maternal bile acid homeostasis has been associated with neurodevelopmental disorders, including ASD, the biological consequences of the specific bile acid changes identified here remain unclear. These findings therefore identify bile acid metabolism as a candidate pathway potentially linking PCB-associated maternal microbiome disturbances to offspring neurodevelopmental alterations.

In contrast, indole-3-propionic acid (IPA), a gut microbiota-derived tryptophan metabolite with reported neuroprotective properties (Owe-Larsson et al. 2025), was significantly decreased (P = 0.0138). Glutathione disulfide (P = 0.0240) and 16,16-dimethyl-prostaglandin D2 (P = 0.0268) were also decreased, suggesting altered glutathione-associated redox metabolism and lipid mediator profiles. However, whether this reflects reduced oxidative stress, enhanced glutathione recycling, or depletion of glutathione reserves remains unclear.

Collectively, these findings demonstrate that maternal PCB FMT exposure alters circulating metabolites associated with bile acid homeostasis, microbial tryptophan metabolism, and redox regulation. The reduction in IPA, alterations in bile acid composition, as well as redox regulation are candidate pathways potentially linking PCB-associated maternal metabolic disturbance to offspring neurodevelopmental alternations. These pathways are known to be important to the gut-brain axis, including modulating host receptor signaling, neuroimmune regulation, and neuronal function (Joyce and O’Malley 2022; Monteiro-Cardoso et al. 2021; Owe-Larsson et al. 2025). However, their causal contributions remain to be established.

### Pathway-level comparison of maternal serum metabolomics and offspring brain transcriptomics identifies candidate pathways linking PCB-associated microbiota to neurodevelopment

Comparing maternal serum metabolomics with P7 offspring brain RNA-seq at the pathway level, as described in Methods, identified several biologically related alterations in microbial metabolite signaling, bile acid metabolism, and oxidative stress-associated pathways (**Figure 9**).

First, decreased maternal IPA coincided with increased expression of *Ahr* (aryl hydrocarbon receptor), Arnt (aryl hydrocarbon receptor nuclear translocator), *Cyp1b1* (cytochrome P450 family 1 subfamily B member 1), and *Nfkb1* (nuclear factor kappa B subunit 1), alongside decreased *Rela* (RELA proto-oncogene, NF-κB subunit) in male offspring brains. These findings suggest altered AhR-associated transcription and NF-κB-related neuroimmune signaling.

Notably, previous experimental evidence demonstrated that microbiota-derived IPA modulates hippocampal AhR/NF-κB signaling, microglial activation, synaptic pruning, and autism-like behaviors, providing mechanistic relevance to our observations (Wang et al. 2023).

Second, increased maternal bile acid concentrations coincided with differential expression of *S1pr2* (sphingosine-1-phosphate receptor 2), *Akt1* (AKT serine/threonine kinase 1), and *Mapk1* (mitogen-activated protein kinase 1) in male offspring brains. Conjugated bile acids can activate S1PR2-mediated AKT and extracellular signal-regulated kinase (ERK) signaling, although this mechanism has been demonstrated primarily in hepatic systems (Kwong et al. 2015; Studer et al. 2012).

Additional changes in *Abcc3* (ATP-binding cassette subfamily C member 3), *Slco1a4* (solute carrier organic anion transporter family member 1A4), and *Cyp27a1* (cytochrome P450 family 27 subfamily A member 1) suggest altered transcription of genes involved in bile acid transport and biosynthesis (Klaassen and Aleksunes 2010). Although these functions have also been primarily characterized in the liver, their differential expression in offspring brain tissue raises the possibility of local alterations in bile acid and sterol metabolism, potentially affecting neurodevelopmental signaling and brain homeostasis.

Third, decreased maternal glutathione disulfide coincided with downregulation of antioxidant-associated genes, including *Gpx4* (glutathione peroxidase 4), *Gpx1* (glutathione peroxidase 1), and *Hmox1* (heme oxygenase 1), in male offspring brains. These findings suggest altered redox-associated transcription, potentially relevant to neuronal vulnerability, as glutathione homeostasis is essential for antioxidant defense and normal brain function and has been implicated in autism spectrum disorder (ASD) (Gu et al. 2015; Ren et al. 2017).

### PCB-FMT alters circulating metabolomic profiles in P7 offspring

To determine whether PCB-FMT influences offspring metabolism, untargeted metabolomic profiling was performed in P7 male and female offspring. Differential metabolite analysis identified three decreased metabolites in males and five altered metabolites in females, including four decreased and one increased metabolite (**Figure 10A and 10B**). These metabolites included triglyceride-, sphingolipid-, phosphatidylcholine-, phosphatidylethanolamine-, and phosphatidylinositol-associated species (**Figure 10C**), suggesting alterations in lipid metabolic homeostasis.

**Figure 8.**
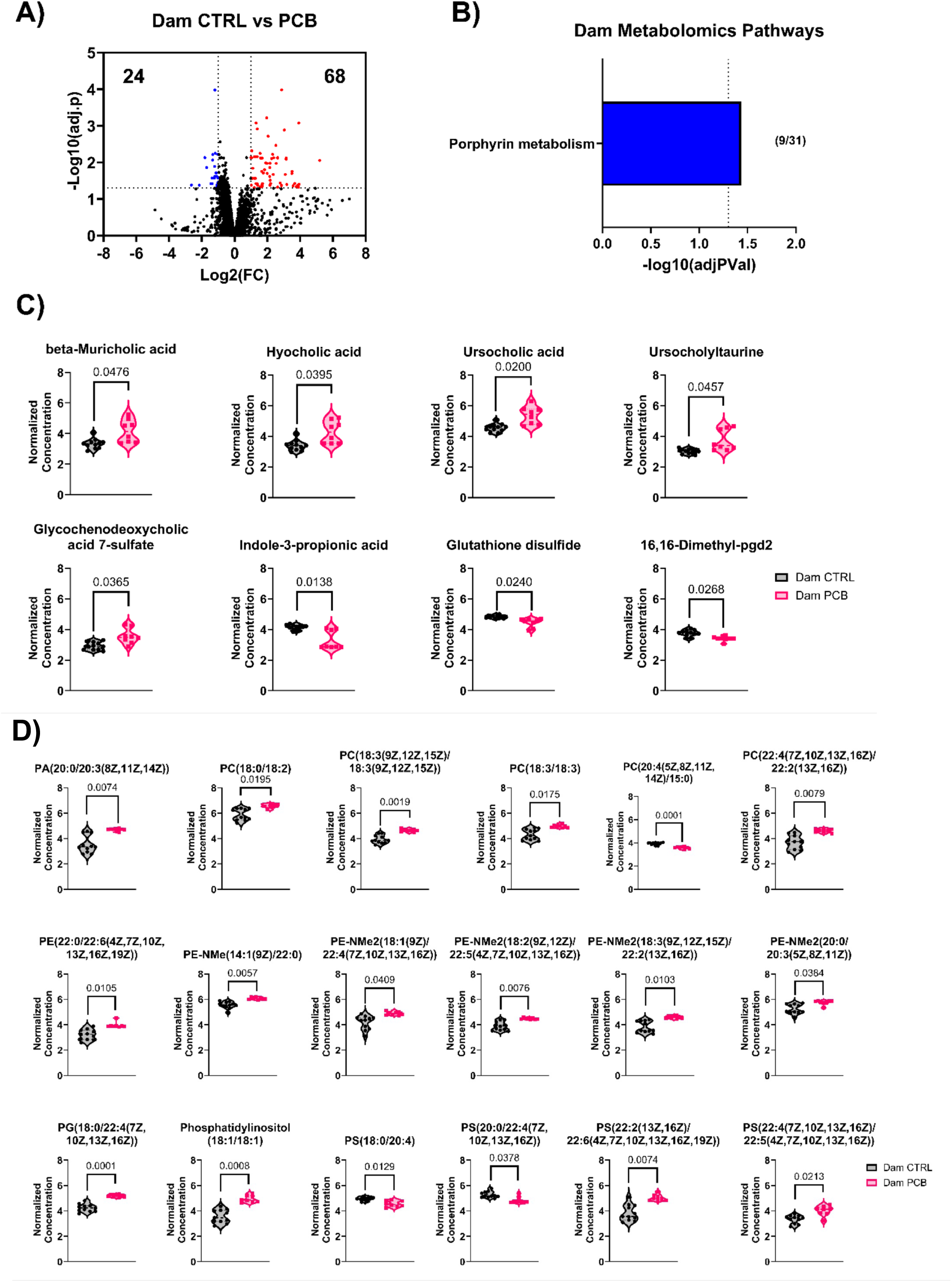
Developmental exposure to the Fox River PCB mixture (FRM) alters maternal serum metabolomic profiles, including bile acid metabolism, microbiome-derived metabolites, redox-associated metabolites, and phospholipid species. (A) Volcano plot showing differential serum metabolite abundance between vehicle-exposed control donor dams (n = 12) and FRM-exposed donor dams (6.0 mg/kg/day; n = 9). Untargeted metabolomic profiling was performed using liquid chromatography–mass spectrometry (LC-MS), and data were analyzed using MetaboAnalyst 6.0 following quantile normalization and log transformation. Black dots represent metabolites not meeting the indicated significance and fold-change thresholds. The x-axis represents log2 fold change, and the y-axis represents −log10(adjusted P value). (B) Metabolic pathway enrichment analysis of significantly altered maternal serum metabolites identified porphyrin metabolism as an enriched pathway, with 9 of 31 pathway-associated metabolites represented. The x-axis indicates −log10(adjusted P value), and the dashed vertical line indicates the nominal P = 0.05 threshold. (C) Selected differentially abundant maternal serum metabolites involved in bile acid metabolism, microbial tryptophan metabolism, redox regulation, and lipid mediator metabolism. Violin plots display normalized metabolite abundance in vehicle-treated controls (black) and FRM-exposed dams (pink), with individual data points shown. P values for individual metabolite comparisons are indicated above the brackets. (D) Selected phospholipid molecular species significantly altered in maternal serum following FRM exposure. Violin plots show normalized metabolite abundance with individual data points. FRM exposure produced lipid species-dependent increases or decreases, consistent with altered circulating phospholipid composition. P values for individual metabolite comparisons are displayed above the brackets. For panels (C) and (D), metabolite abundance was compared between vehicle-exposed and FRM-exposed donor dams (n = 12 and n = 9, respectively). P values are displayed as provided by the metabolomics analysis.

**Figure 9.**
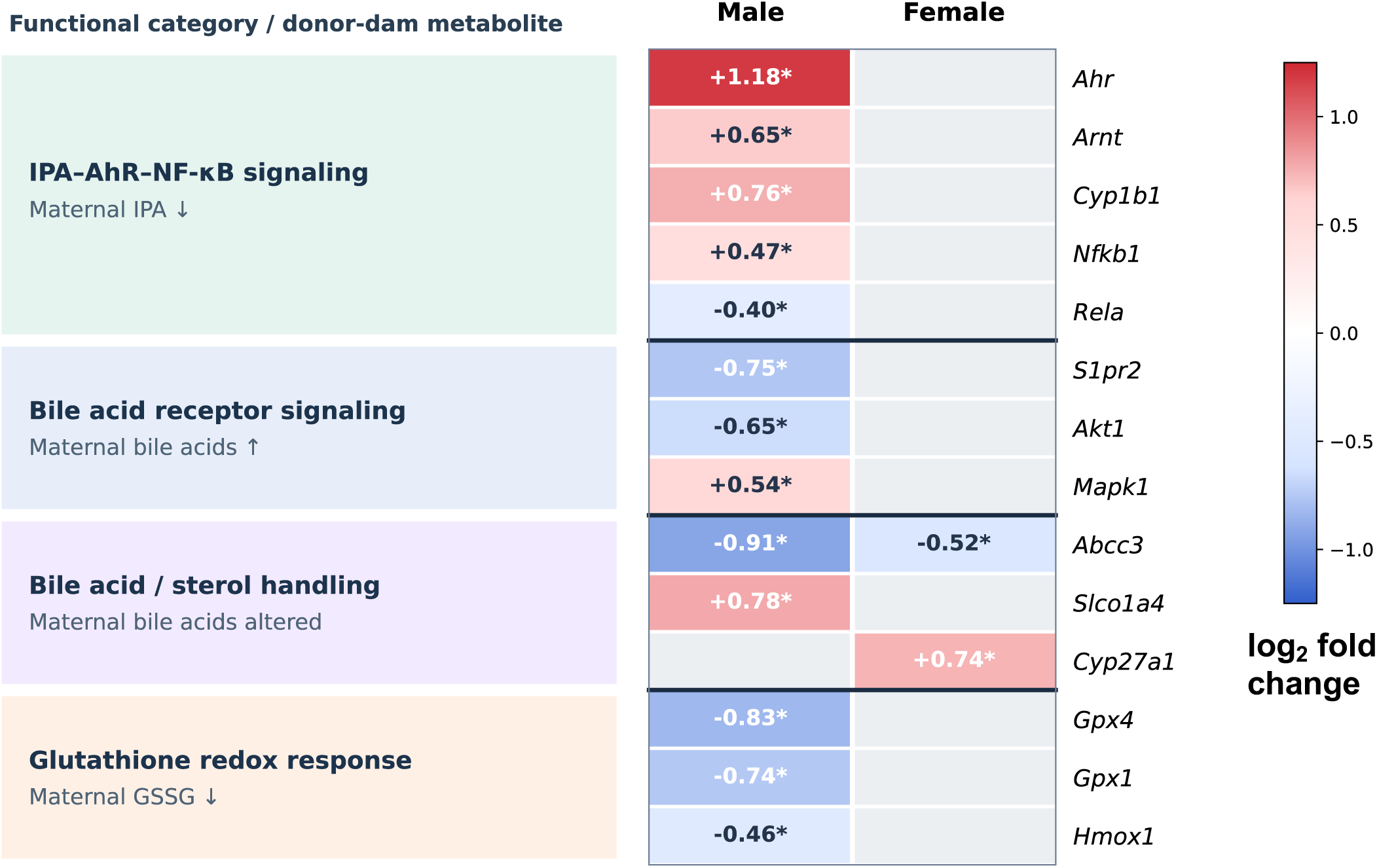
Pathway-level comparison of maternal serum metabolomics and offspring brain transcriptomics identifies candidate pathways linking PCB-associated gut microbiota to neurodevelopment. Heatmap showing log2 fold changes of 14 selected genes in P7 offspring brains categorized by biological functions associated with maternal metabolic alterations, including IPA-AhR-NF-κB signaling, bile acid receptor signaling, bile acid and sterol handling, and glutathione redox regulation. Whole-brain RNA-seq was performed in P7 offspring (n = 4 per sex per FMT group; total n = 16). Differential expression was analyzed using limma-voom with sample quality weights and FDR-BH < 0.05. Red and blue indicate upregulation and downregulation, respectively. Asterisks (*) indicate statistically significant differential expression, while gray cells indicate unavailable log2 fold-change values from the significant-DEG datasets. Candidate genes were selected from pathways functionally associated with the altered maternal metabolites, as described in Methods, rather than by statistical integration.

**Figure 10.**
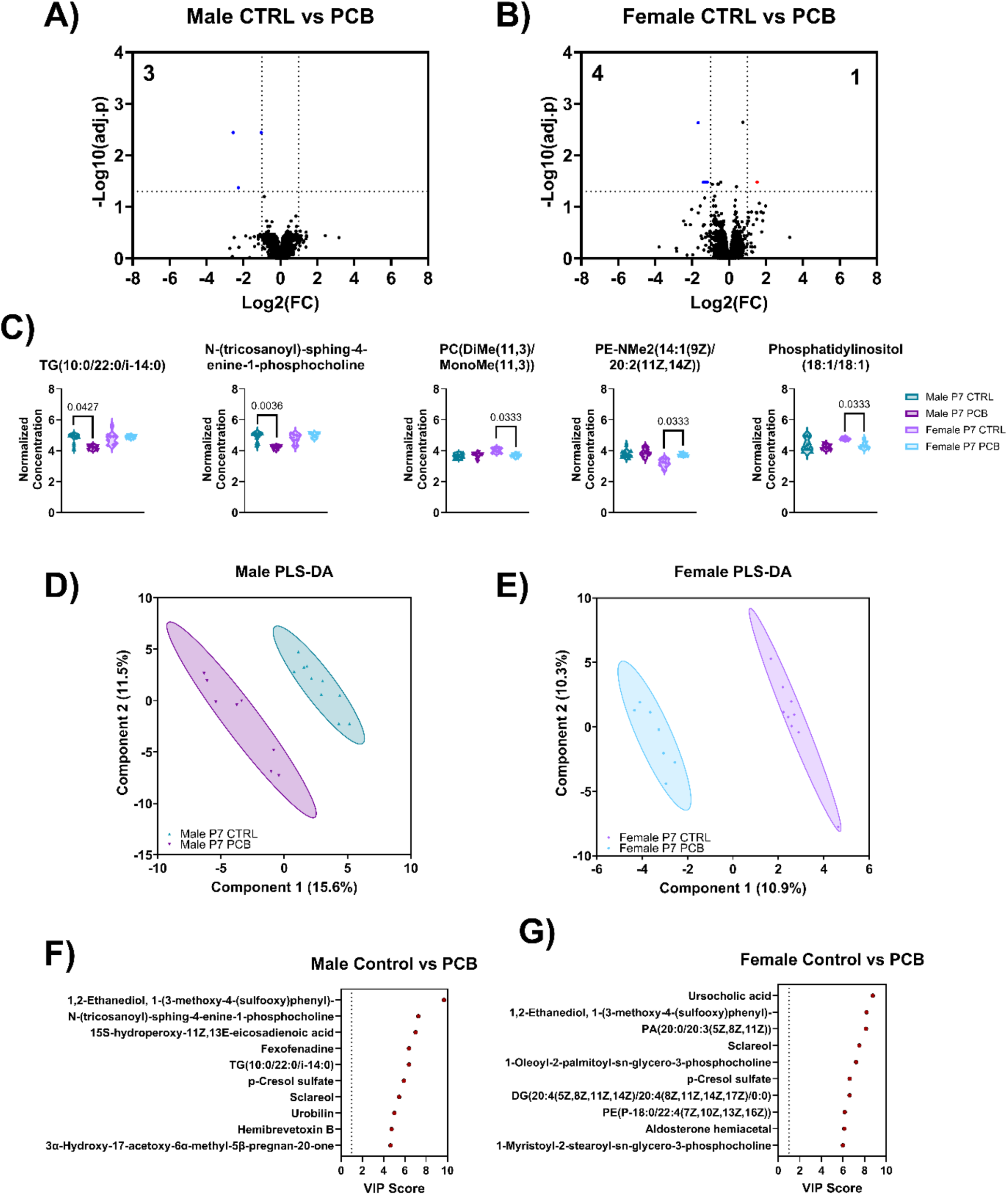
PCB-associated FMT alters circulating metabolomic profiles in P7 male and female offspring. (A-B) Volcano plots showing differential serum metabolite abundance in male (**A**) and female (**B**) P7 offspring following PCB-associated FMT compared with vehicle-associated FMT. Untargeted serum metabolomics was performed in male controls (n = 11), male PCB-FMT (n = 9), female controls (n = 10), and female PCB-FMT (n = 8). Metabolomic data were quantile-normalized, log-transformed, and analyzed using MetaboAnalyst 6.0. Red and blue dots indicate significantly increased and decreased metabolites, respectively, whereas black dots represent metabolites not meeting the indicated significance and fold-change thresholds. The x-axis represents log2 fold change, and the y-axis represents −log10(adjusted P value). **(C)** Selected differentially abundant serum metabolites associated with triglyceride, sphingolipid, phosphatidylcholine, phosphatidylethanolamine, and phosphatidylinositol metabolism. Violin plots show normalized metabolite abundance with individual data points for male and female offspring in the vehicle-FMT and PCB-associated FMT groups. P values for individual comparisons are displayed above the brackets. **(D-E)** Partial least squares discriminant analysis (PLS-DA) of serum metabolomic profiles in male (D) and female (E) P7 offspring. Each point represents an individual biological sample, with colors distinguishing vehicle-FMT and PCB-associated FMT groups. Ellipses illustrate group distributions, and the percentages on the axes indicate variance represented by the respective PLS-DA components. **(F-G)** Variable importance in projection (VIP) analysis identifying the top 10 metabolites contributing to PLS-DA group discrimination in male (**F**) and female (**G**) offspring. Metabolites are ranked according to their VIP scores, with higher scores indicating greater contributions to group separation.

**Figure 11.**
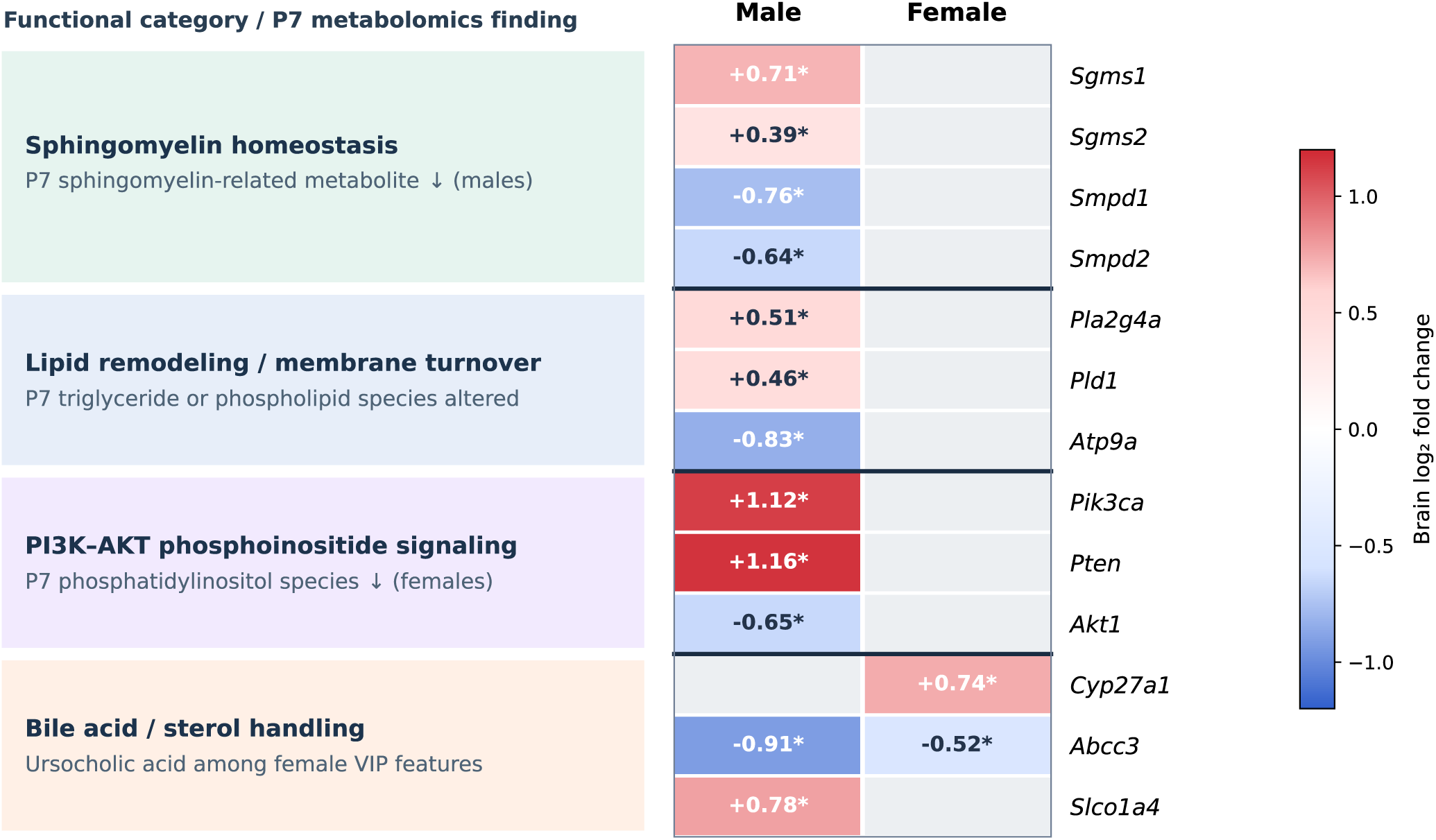
Pathway-level comparison of P7 offspring serum metabolomics and brain transcriptomics identified candidate pathways linking PCB-associated gut microbiota to neurodevelopmental alterations. Heatmap showing log2 fold changes of 13 selected genes categorized by biological functions associated with offspring serum metabolic alterations, including sphingomyelin homeostasis, lipid remodeling and membrane turnover, PI3K-AKT phosphoinositide signaling, and bile acid and sterol handling. Untargeted serum metabolomics was performed in P7 offspring (male vehicle-FMT, n = 11; male PCB-FMT, n = 9; female vehicle-FMT, n = 10; female PCB-FMT, n = 8). Whole-brain RNA sequencing was performed using n = 4 biological replicates per sex per FMT group (total n = 16). Differential gene expression was analyzed separately by sex using limma-voom with sample quality weights and FDR-BH < 0.05. Red and blue indicate upregulation and downregulation, respectively, with numerical log2 fold changes displayed within cells. Asterisks (*) indicate statistically significant differential expression (FDR-BH < 0.05). Metabolite-gene correspondences were curated from established biological pathway relationships, as described in Methods, rather than derived statistically.

Partial least squares discriminant analysis (PLS-DA) showed separation between vehicle-FMT and PCB-FMT groups in both males and females (**Figure 10D and 10E**). Variable importance in projection (VIP) analysis identified lipid-associated metabolites, p-cresol sulfate, and other metabolic features as major contributors to group discrimination (**Figure 10F and 10G**). Notably, ursocholic acid, which was elevated in PCB-exposed donor dams **(Figure 8**), was also identified as an important discriminating feature in female offspring.

### Pathway-level comparison of P7 offspring serum metabolomics and brain transcriptomics identifies candidate lipid metabolism and signaling pathways

To identify potential molecular connections between circulating metabolic alterations and brain transcriptional responses following PCB-FMT, we compared P7 offspring serum metabolomics with whole-brain RNA-seq data at the pathway level, as described in Methods (**Figure 11**). Because only three metabolites in male and five in female offspring were significantly altered, this comparison is exploratory. In male offspring, decreased circulating abundance of a sphingomyelin-related metabolite coincided with increased brain expression of sphingomyelin synthase genes *Sgms1* (log2FC = 0.71) and *Sgms2* (log2FC = 0.39), alongside decreased expression of sphingomyelinase genes *Smpd1* (log2FC = -0.76) and *Smpd2* (log2FC = -0.64), suggesting altered sphingomyelin homeostasis. Additional male-associated transcriptional alterations included increased expression of *Pla2g4a* (log2FC = 0.51) and *Pld1* (log2FC = 0.46), together with decreased *Atp9a* (log2FC = -0.83), indicating changes in genes involved in phospholipid remodeling and membrane trafficking. Furthermore, *Pik3ca* (log2FC = 1.12) and *Pten* (log2FC = 1.16) were upregulated, whereas *Akt1* (log2FC = -0.65) was downregulated in male brains, suggesting altered transcriptional regulation of PI3K-AKT-associated signaling. This correspondence is reported with the caveat that the associated phosphatidylinositol alterations were detected in female rather than male serum; the tow observations are therefore not matched within sex and are presented only as a shared pathway-level theme. Notably, this pathway-level association involved phosphatidylinositol alterations detected in female serum rather than male serum. Bile acid- and sterol-associated genes also exhibited differential expression, including decreased *Abcc3* in both males (log2FC = -0.91) and females (log2FC = - 0.52), increased *Slco1a4* in males (log2FC = 0.78), and increased *Cyp27a1* in females (log2FC = 0.74). These transcriptional alterations were considered alongside the identification of ursocholic acid as a discriminating metabolomic feature in female offspring.

## Discussion

Taken together, this study demonstrated that transplanting gut microbiota from dams exposed to the environmentally relevant FRM reproduced early-life neurobehavioral alterations and induced extensive molecular changes in offspring without direct PCB administration to recipient animals (**Figure 12**). PCB-FMT significantly reduced ultrasonic vocalizations in P7 offspring, accompanied by pronounced sex-dependent brain transcriptomic responses. Both sexes exhibited downregulation of genes associated with neurotransmitter receptor signaling and ion channel function, alongside upregulation of RNA processing pathways. Male offspring displayed more extensive transcriptional alterations, characterized by downregulation of protein synthesis-related pathways and upregulation of RHO GTPase signaling and neuronal regulatory pathways, whereas female offspring exhibited alterations involving GPCR signaling, DNA repair-associated pathways, and extracellular matrix remodeling. Untargeted serum metabolomics of PCB-exposed donor dams revealed increased concentrations of selected bile acids, decreased the microbial tryptophan metabolite IPA and glutathione disulfide, and altered phospholipid profiles. Metabolomic analysis of P7 offspring further identified sex-associated alterations in sphingolipid, phospholipid, and bile acid-related metabolites. Integrating metabolomic and transcriptomic findings highlighted IPA-AhR-NFκB signaling, bile acid receptor signaling and sterol metabolism, phospholipid remodeling, and redox regulation as candidate pathways potentially linking PCB-associated gut microbiota to altered neurodevelopmental programming. Together, our findings support the role of PCB-associated gut microbiota in developmental neurotoxicity and identify candidate metabolic and molecular mechanisms that warrant further investigation. Although the integrated pathways remain hypothesis-generating and potential residual PCB transfer through donor inocula has not been excluded, this study provides a framework for understanding how environmental pollutant-associated changes in the maternal gut microbiome may influence offspring brain development and neurobehavioral outcomes.

**Figure 12.**
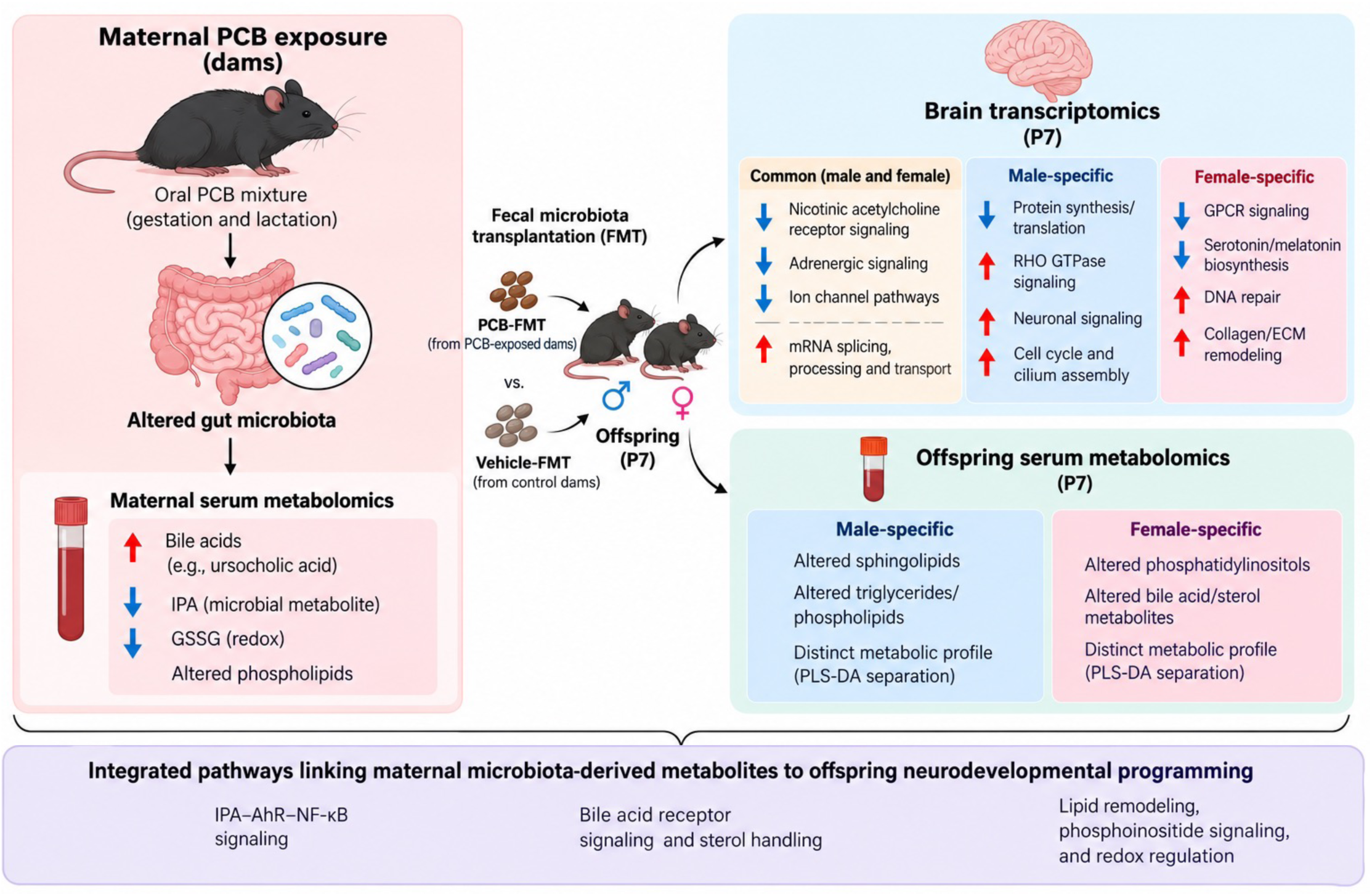
Schematic summary of maternal PCB-associated gut microbiota-mediated alterations in offspring neurodevelopment. FMT from dams exposed to the environmentally relevant Fox River PCB mixture reduced ultrasonic vocalizations in P7 offspring and induced sex-dependent alterations in brain transcriptomic and serum metabolomic profiles. Both sexes exhibited suppression of neurotransmitter signaling and ion channel pathways, as well as increased RNA processing pathways. Male and female offspring displayed distinct changes in neuronal signaling, protein synthesis, DNA repair, and extracellular matrix remodeling. Maternal serum metabolomics revealed alterations in bile acids, indole-3 propionic acid, glutathione disulfide, and phospholipids, while offspring exhibited sex-specific changes in lipid metabolism. Integrated multi-omics analyses identified IPA-AhR-NF-κB signaling, bile acid receptor signaling, phospholipid remodeling, and redox regulation as candidate mechanisms linking PCB-associated maternal gut microbiota to altered offspring neurodevelopment. These proposed pathways are hypothesis-generating and require further mechanistic validation. Disclaimer: Generative AI was used only for figure graphics, not for scientific ideas, experimental design, or data.

The environmentally relevant FRM was selected to approximate the complex PCB congener profile associated with contaminated fish from the Fox River/Green Bay region. The exposure regimen used here has been characterized previously with respect to both internal PCB burden and neurobehavioral outcome (Li et al. 2024; Wilson et al. 2024; Yang et al. 2009), and exposure relevance is better judged from internal burdens and congener-specific profiles than from administered dose.

A major finding of this study is that transplantation of intestinal microbiota-associated material from PCB-exposed dams reproduced the reduction in neonatal ultrasonic vocalizations previously observed following direct maternal PCB exposure (Wilson et al. 2024). Because recipient dams were not directly administered PCBs, these findings support the hypothesis that PCB-associated alterations in the intestinal microbial ecosystem contribute to developmental neurobehavioral toxicity. This interpretation is consistent with previous studies demonstrating that transplantation of disease-associated gut microbiota can modify recipient neurological and behavioral phenotypes (Sharon et al. 2019), and with evidence that developmental exposure to a human-relevant PCB mixture in the maternal diet disrupts host-microbe interactions in weanling offspring (Rude et al. 2019). Nevertheless, fecal transplantation transfers a complex mixture of microorganisms and associated metabolites, and potential residual PCB contamination of the donor inocula cannot be excluded. Consequently, the present findings support a transferable PCB-associated neurobehavioral phenotype but do not yet establish that microbiota composition alone is sufficient to produce these effects.

Although PCB-associated FMT induced extensive transcriptional alterations in offspring brains, changes in mRNA abundance do not necessarily correspond to proportional changes in protein abundance or biological activity. Protein expression is additionally regulated by RNA stability, translation efficiency, post-translational modifications, and protein degradation (Buccitelli and Selbach 2020; Vogel and Marcotte 2012). This distinction is particularly relevant to the enrichment of translation-associated pathways among downregulated genes in male offspring, which indicates altered transcription of protein synthesis-related machinery but does not directly demonstrate reduced global protein synthesis. Similarly, increased expression of genes associated with neuronal signaling, RHO GTPase regulation, or RNA processing does not necessarily indicate increased pathway activity. Furthermore, because RNA-seq was performed using whole-brain homogenates, regional or cell-type-specific transcriptional changes may have been masked or influenced by differences in cellular composition. Future studies integrating quantitative proteomics, ribosome profiling, and region- or cell-type-resolved transcriptomics will be essential to determine the functional consequences of these molecular responses in the developing brain.

An intriguing finding was the marked difference in the magnitude of brain transcriptional responses between male and female offspring despite the absence of a significant sex-dependent effect on neonatal ultrasonic vocalizations. Male offspring exhibited substantially more differentially expressed genes than females, with prominent alterations in protein synthesis, RHO GTPase signaling, and neuronal regulatory pathways, whereas female offspring exhibited transcriptional changes associated with GPCR signaling, DNA repair-related processes, and extracellular matrix remodeling. Nevertheless, both sexes shared downregulation of genes involved in neurotransmitter signaling and ion channel function, suggesting potentially convergent neurodevelopmental effects despite distinct broader molecular responses. Furthermore, neonatal ultrasonic vocalizations represent a developmentally restricted behavioral endpoint and may not capture molecular alterations that manifest as sex-dependent phenotypes later in life. Previous studies have demonstrated substantial developmental and individual variability in mouse pup ultrasonic vocalizations (Rieger and Dougherty 2016; Yin et al. 2016). Together, these observations suggest that male and female offspring may respond to PCB-associated microbiota through distinct molecular routes that converge on a comparable adverse outcome, and that the sex-opposing regulation of inflammation-resolution, extracellular matrix, and neurite development pathways may be particularly consequential for early social communication, of which isolation-induced ultrasonic vocalization is the earliest measurable index. Sex-biased molecular responses may therefore precede or occur independently of detectable sex differences in early vocal communication.

The PCB-altered serum metabolites in dams were in classes tied to gut microbial activity. Elevated circulating beta-muricholic acid reproduces what we previously observed in female mice following oral exposure to the FRM (Cheng et al. 2018), supporting the reliability of the bile acid signal in this model. Multiple bile acids were significantly elevated following FRM exposure, including beta-muricholic acid, hyocholic acid, ursocholic acid, and their conjugated and sulfated derivatives ursocholyltaurine and glycochenodeoxycholic acid 7-sulfate. Because the gut microbiome mediates the deconjugation, dehydroxylation, and sulfation reactions that convert primary bile acids into secondary and microbially modified species (Collins et al. 2023), the coordinated elevation of these bile acids is consistent with altered microbial bile acid metabolism. Bile acids additionally act as signaling ligands through host receptors including FXR and TGR5 (Chiang and Ferrell 2020), pathways with recognized roles in the lipid and energy dysregulation associated with PCB toxicity (Wahlang et al. 2019). Other bile acids, such as hyocholic acid and ursocholic acid, represent previously uninvestigated opportunities to link microbial bile acid metabolism to PCB exposure.

Several metabolites involved in inflammation and intestinal barrier function were altered following FRM exposure. IPA, a metabolite produced by gut bacteria that supports intestinal barrier integrity (Li et al. 2025) was significantly decreased. Additionally, the prostaglandin derivative 16,16-dimethyl-PGD2 and glutathione disulfide were decreased, consistent with altered inflammatory and oxidative-stress signaling (Guan 2023; Joo and Sadikot 2012). Overall, this reduction indicates that the PCB-associated microbiome has a potentially less protective metabolic profile.

Circulating phospholipids were broadly altered following FRM exposure, with most increasing in abundance. Phospholipids are major components of circulating lipoproteins, where they transport triglycerides, cholesterol, and fatty acids between tissues (Ginsberg 1998). Among the altered metabolites were multiple phosphatidylcholine (PC), phosphatidylethanolamine (PE), phosphatidylinositol (PI), phosphatidylserine (PS), phosphatidylglycerol (PG), and phosphatidic acid (PA) species, suggesting widespread remodeling of glycerophospholipid (Bargui et al. 2021). PCB-associated alterations in circulating phospholipid metabolism have been reported previously, including increased total serum phospholipids and redistribution of phospholipids among lipoprotein fractions in PCB-exposed rats (Kato et al. 1982), altered serum glycerophospholipid profiles following dietary Aroclor 1260 exposure in pigs (Hernández-Mesa et al. 2022), and PCB-associated changes in the polyunsaturated fatty acid composition of serum phospholipids in human populations (Grandjean and Weihe 2003). Together, the increased circulating phospholipids suggest altered phospholipid synthesis and remodeling.

KEGG pathway analysis identified a single significantly altered pathway, porphyrin metabolism, for the dam metabolomics. Porphyrin metabolism includes the biosynthesis and degradation of heme and related tetrapyrroles, which are essential cofactors for oxygen transport, mitochondrial electron transport, and cytochrome P450–mediated xenobiotic metabolism (Phillips 2019). Alteration of porphyrin metabolism following FRM exposure is consistent with previous evidence that PCBs disrupt heme biosynthesis and porphyrin homeostasis. Aroclor 1254 exposure has been shown to increase circulating protoporphyrin in rats in a dose- and time-dependent manner (Cho et al. 2003), while additional studies have demonstrated hepatic accumulation and increased urinary excretion of porphyrins following PCB exposure in rodents and humans (Chang et al. 1980; Goldstein et al. 1974). Together, these findings suggest that porphyrin metabolism represents a conserved metabolic target of direct PCB exposure in dams; whether PCB-associated alterations in the gut microbiome are sufficient to influence this pathway in the absence of direct exposure was not tested here.

Interestingly, offspring of PCB-exposed FMT dams exhibited substantially fewer metabolic alterations than those observed in the dams. No differences were observed between male and female offspring; however, a small number of metabolites differed between maternal FMT exposure groups. In male pups, pinonic acid, TG(10:0/22:0/i-14:0), and N-(tricosanoyl)-sphing-4-enine-1-phosphocholine were decreased. TG(10:0/22:0/i-14:0) is a triacylglycerol, which is involved in fatty acid storage and lipid homeostasis (Klemm and Carvalho 2024). Although this specific triacylglycerol has not been previously associated with PCBs, PCB exposure has been linked to disrupted triglyceride metabolism. This finding is consistent with male Sprague-Dawley rats exposed to PCB126, in which triglyceride levels decreased over time (Gadupudi et al. 2016). In contrast, short-term exposure to PCB118, PCB153, or a mixture of PCB118 and PCB153 increased circulating triglycerides (Mesnier et al. 2015). N-(Tricosanoyl)-sphing-4-enine-1-phosphocholine, also named SM(d18:1/23:0), is a sphingomyelin, which contributes to cellular membrane structure (Ramstedt and Slotte 2002). Although PCB exposure has been associated with disruption of sphingolipid metabolism in hepatic and cellular models (Deng et al. 2019; Mesnage et al. 2018), effects on circulating sphingomyelins appear less consistent. PCB126 alone did not significantly alter total plasma sphingomyelin levels in mice (Deng et al. 2020), whereas exposure to a broader persistent organic pollutant mixture containing PCBs was associated with decreased circulating sphingomyelins (Sinioja et al. 2022). Together, these findings suggest that maternal transfer of an FRM-associated gut microbiome produces only minor metabolic effects in male offspring, with the limited alterations primarily involving lipids associated with triglyceride and sphingolipid metabolism.

Female offspring similarly exhibited only limited metabolic changes. PE-NMe2(14:1(9Z)/20:2(11Z,14Z)) was increased, whereas 1-hexadecanoyl-2-(9Z-octadecenoyl)-sn-glycero-3-phospho-(1’-myo-inositol), phosphatidylinositol (18:1/18:1), PC(DiMe(11,3)/MonoMe (11,3)), and panaxytriol were decreased. As in dams, altered PE, PI, and PC suggest subtle phospholipid synthesis and remodeling, though to a lesser extent. PLS-DA further supported the changes observed in female offspring, with several PC, PE, and PA metabolites contributing most strongly to group separation. Notably, ursocholic acid was the highest-ranking VIP metabolite in female offspring. Interestingly, ursocholic acid was significantly elevated in FMT dams, but decreased in female offspring, suggesting that maternal transfer of a PCB-associated microbiome may influence bile acid metabolism across generations. Together, these findings suggest that although female offspring exhibited fewer metabolic alterations than the FMT dams, their metabolic profile more closely paralleled the dams than that of male offspring, particularly phospholipid and bile acid-associated changes.

Although the present study identified neurobehavioral, transcriptomic, and metabolomic alterations following transplantation of PCB-associated intestinal microbiota, microbial community composition and functional potential were not directly characterized. The primary objective was to determine whether transfer of PCB-associated intestinal material could reproduce aspects of developmental neurobehavioral toxicity and to identify corresponding molecular changes in offspring. Previous microbiota transplantation studies have demonstrated the value of combining microbial community profiling with metabolomics and host molecular analyses to identify candidate mechanisms linking gut microbiota to neurobehavioral outcomes (Sharon et al. 2019). In the present study, decreased IPA and altered secondary bile acid profiles in PCB-exposed donor dams, together with changes in offspring circulating lipids and brain transcriptional pathways, highlight candidate microbiome-associated metabolic processes for further investigation. Future studies integrating shotgun metagenomic sequencing, microbial functional profiling, targeted metabolomics, and microbial engraftment analysis in donor dams, FMT recipients, and their offspring will be necessary to establish direct microbiome–metabolite–brain relationships. In addition, independently prepared donor inocula and defined microbial communities will help distinguish microbiota-dependent effects from contributions of nonmicrobial components of the transplanted material. Because large intestinal contents from all donor dams within an exposure group were pooled into a single inoculum, each group was represented by one donor community; donor-to-donor variability is therefore not captured, and recipient outcomes cannot be attributed to individual donor microbiomes. Finally, because untargeted metabolomics was performed in directly exposed donor dams whereas RNA sequencing was performed in offspring of FMT recipients, there pathway-level correspondences described here identify candidate relationships rather than demonstrated metabolic transmission, and establishing causality will require metabolomic and transcriptomic measurements in the same animals.

More broadly, our findings highlight the importance of considering the gut microbiome as an integral component of the biological response to environmental toxicants, rather than simply a secondary target of chemical exposure. Developmental neurotoxicity has traditionally been investigated through the direct effects of environmental chemicals on the developing nervous system. Our findings suggest that PCB-associated alterations in the maternal intestinal environment may represent an additional route through which environmental exposures influence offspring neurodevelopment. The pronounced sex-biased brain transcriptional responses, together with shared neurobehavioral alterations, further underscore the complexity of microbiome–brain interactions during early development. Although the specific microbial mediators and causal metabolic pathways remain to be established, these observations support a broader conceptual framework in which environmental exposures, gut microbial metabolism, and host molecular responses collectively shape neurodevelopmental susceptibility. Understanding these interactions may ultimately reveal previously unrecognized mechanisms of developmental neurotoxicity and identify opportunities to mitigate the long-term neurological consequences of early-life environmental exposures to PCB contaminants.

## Acknowledgements

The authors would like to thank members of the Cui, Lehmler, and Lein Laboratories for proof-reading the manuscript.

## Footnote

This work was supported by the National Institute of Environmental Health Sciences (NIEHS), National Institutes of Health (NIH), through grants R01ES031098, R01ES030197, and P30ES030287, Environmental Health and Microbiome Research Center (EHMBRACE) at University of Washington (Cui), as well as startup funding (Cui) from the Department of Environmental and Molecular Toxicology, College of Agricultural Sciences, Oregon State University.

## References

1. Anezaki, K., & Nakano, T. (2014). Concentration levels and congener profiles of polychlorinated biphenyls, pentachlorobenzene, and hexachlorobenzene in commercial pigments. Environmental science and pollution research international, 21(2), 998–1009.

2. Annamneedi A, Gora C, Dudas A, Leray X, Bozon V, Crepieux P, Pellissier LP. 2025. Towards the convergent therapeutic potential of g protein-coupled receptors in autism spectrum disorders. Br J Pharmacol. 182(14):3044–3067.

3. Bargui R, Solgadi A, Prost B, Chester M, Ferreiro A, Piquereau J, Moulin M. 2021. Phospholipids: Identification and implication in muscle pathophysiology. Int J Mol Sci. 22(15).

4. Basil MC, Levy BD. 2016. Specialized pro-resolving mediators: Endogenous regulators of infection and inflammation. Nat Rev Immunol. 16(1):51–67.

5. Baxter AJ, Brugha TS, Erskine HE, Scheurer RW, Vos T, Scott JG. 2015. The epidemiology and global burden of autism spectrum disorders. Psychol Med. 45(3):601–613.

6. Buccitelli C, Selbach M. 2020. Mrnas, proteins and the emerging principles of gene expression control. Nat Rev Genet. 21(10):630–644.

7. Chang KJ, Lu FJ, Tung TC, Lee TP. 1980. Studies on patients with polychlorinated biphenyl poisoning. 2. Determination of urinary coproporphyrin, uroporphyrin, delta-aminolevulinic acid and porphobilinogen. Res Commun Chem Pathol Pharmacol. 30(3):547–554.

8. Cheng SL, Li X, Lehmler H-J, Phillips B, Shen D, Cui JY. 2018. Gut microbiota modulates interactions between polychlorinated biphenyls and bile acid homeostasis. Toxicol Sci. 166(2):269–287.

9. Chiang JYL, Ferrell JM. 2020. Bile acid receptors fxr and tgr5 signaling in fatty liver diseases and therapy. American Journal of Physiology-Gastrointestinal and Liver Physiology. 318(3):G554–G573.

10. Cho JH, Jeong SH, Yun HI. 2003. Changes of urinary and blood porphyrin profiles by exposure to pcbs, lead or diazinon in rats. Vet Hum Toxicol. 45(4):193–198.

11. Collins SL, Stine JG, Bisanz JE, Okafor CD, Patterson AD. 2023. Bile acids and the gut microbiota: Metabolic interactions and impacts on disease. Nature Reviews Microbiology. 21(4):236–247.

12. Deng P, Barney J, Petriello MC, Morris AJ, Wahlang B, Hennig B. 2019. Hepatic metabolomics reveals that liver injury increases pcb 126-induced oxidative stress and metabolic dysfunction. Chemosphere. 217:140–149.

13. Deng P, Wang C, Wahlang B, Sexton T, Morris AJ, Hennig B. 2020. Co-exposure to pcb126 and pfos increases biomarkers associated with cardiovascular disease risk and liver injury in mice. Toxicology and Applied Pharmacology. 409:115301.

14. Doi, H., Furui, A., Ueda, R., Shimatani, K., Yamamoto, M., Eguchi, A., Sagara, N., Sakurai, K., Mori, C., & Tsuji, T. (2024). Risk of autism spectrum disorder at 18 months of age is associated with prenatal level of polychlorinated biphenyls exposure in a Japanese birth cohort. Scientific reports, 14(1), 31872.

15. Dreiem A, Okoniewski RJ, Brosch KO, Miller VM, Seegal RF. 2010. Polychlorinated biphenyls and polybrominated diphenyl ethers alter striatal dopamine neurochemistry in synaptosomes from developing rats in an additive manner. Toxicol Sci. 118(1):150–159.

16. Duncan BW, Murphy KE, Maness PF. 2021. Molecular mechanisms of l1 and ncam adhesion molecules in synaptic pruning, plasticity, and stabilization. Front Cell Dev Biol. 9:625340.

17. Elangovan S, Borody TJ, Holsinger RMD. 2022. Fecal microbiota transplantation reduces pathology and improves cognition in a mouse model of alzheimer’s disease. Cells. 12(1).

18. Falace A, Nosrati MSS, Dostmohammadi A, Doustmohammadi M, Striano P, Zara F, Hamada N, Nagata KI, Scala M. 2026. Wiring the developing brain: The rho gtpase signaling network in neurodevelopmental disorders. Neurobiol Dis. 229:107609.

19. Gadupudi GS, Klaren WD, Olivier AK, Klingelhutz AJ, Robertson LW. 2016. Pcb126-induced disruption in gluconeogenesis and fatty acid oxidation precedes fatty liver in male rats. Toxicol Sci. 149(1):98–110.

20. Gilbert ME, Crofton KM. 1999. Developmental exposure to a commercial pcb mixture (aroclor 1254) produces a persistent impairment in long-term potentiation in the rat dentate gyrus in vivo. Brain Res. 850(1-2):87–95.

21. Ginsberg HN. 1998. Lipoprotein physiology. Endocrinol Metab Clin North Am. 27(3):503–519.

22. Goldstein JA, Hickman P, Jue DL. 1974. Experimental hepatic porphyria induced by polychlorinated biphenyls. Toxicology and Applied Pharmacology. 27(2):437–448.

23. Goo N, Bae HJ, Park K, Kim J, Jeong Y, Cai M, Cho K, Jung SY, Kim DH, Ryu JH. 2020. The effect of fecal microbiota transplantation on autistic-like behaviors in fmr1 ko mice. Life Sci. 262:118497.

24. Grabrucker S, Marizzoni M, Silajdzic E, Lopizzo N, Mombelli E, Nicolas S, Dohm-Hansen S, Scassellati C, Moretti DV, Rosa M et al. 2023. Microbiota from alzheimer’s patients induce deficits in cognition and hippocampal neurogenesis. Brain. 146(12):4916–4934.

25. Grandjean P, Weihe P. 2003. Arachidonic acid status during pregnancy is associated with polychlorinated biphenyl exposure123. The American Journal of Clinical Nutrition. 77(3):715–719.

26. Gu F, Chauhan V, Chauhan A. 2015. Glutathione redox imbalance in brain disorders. Curr Opin Clin Nutr Metab Care. 18(1):89–95.

27. Guan X. 2023. Glutathione and glutathione disulfide – their biomedical and pharmaceutical applications. Med Chem Res. 32(9):1972–1994.

28. Guo D, Yang X, Shi L. 2020. Rho gtpase regulators and effectors in autism spectrum disorders: Animal models and insights for therapeutics. Cells. 9(4).

29. Hahn ME, Lavooy MJ. 2005. A review of the methods of studies on infant ultrasound production and maternal retrieval in small rodents. Behav Genet. 35(1):31–52.

30. Hazan S. 2020. Rapid improvement in alzheimer’s disease symptoms following fecal microbiota transplantation: A case report. J Int Med Res. 48(6):300060520925930.

31. Herkert, N. J., Jahnke, J. C., & Hornbuckle, K. C. (2018). Emissions of Tetrachlorobiphenyls (PCBs 47, 51, and 68) from Polymer Resin on Kitchen Cabinets as a Non-Aroclor Source to Residential Air. Environmental science & technology, 52(9), 5154–5160.

32. Hernández-Mesa M, Narduzzi L, Ouzia S, Soetart N, Jaillardon L, Guitton Y, Le Bizec B, Dervilly G. 2022. Metabolomics and lipidomics to identify biomarkers of effect related to exposure to non-dioxin-like polychlorinated biphenyls in pigs. Chemosphere. 296:133957.

33. Hu, D., & Hornbuckle, K. C. (2010). Inadvertent polychlorinated biphenyls in commercial paint pigments. Environmental science & technology, 44(8), 2822–2827.

34. Jacobson JL, Jacobson SW. 1996. Intellectual impairment in children exposed to polychlorinated biphenyls in utero. N Engl J Med. 335(11):783–789.

35. Jacobson JL, Jacobson SW. 2003. Prenatal exposure to polychlorinated biphenyls and attention at school age. J Pediatr. 143(6):780–788.

36. Jang HM, Kim JK, Joo MK, Shin YJ, Lee CK, Kim HJ, Kim DH. 2021. Transplantation of fecal microbiota from patients with inflammatory bowel disease and depression alters immune response and behavior in recipient mice. Sci Rep. 11(1):20406.

37. Joo M, Sadikot RT. 2012. Pgd synthase and pgd2 in immune resposne. Mediators Inflamm. 2012:503128.

38. Joo Y, Benavides DR. 2021. Local protein translation and rna processing of synaptic proteins in autism spectrum disorder. Int J Mol Sci. 22(6).

39. Joyce SA, O’Malley D. 2022. Bile acids, bioactive signalling molecules in interoceptive gut-to-brain communication. J Physiol. 600(11):2565–2578.

40. Kang DW, Adams JB, Coleman DM, Pollard EL, Maldonado J, McDonough-Means S, Caporaso JG, Krajmalnik-Brown R. 2019. Long-term benefit of microbiota transfer therapy on autism symptoms and gut microbiota. Sci Rep. 9(1):5821.

41. Kang DW, Adams JB, Gregory AC, Borody T, Chittick L, Fasano A, Khoruts A, Geis E, Maldonado J, McDonough-Means S et al. 2017. Microbiota transfer therapy alters gut ecosystem and improves gastrointestinal and autism symptoms: An open-label study. Microbiome. 5(1):10.

42. Kato N, Kawai K, Yoshida A. 1982. Effects of dietary polychlorinated biphenyls and protein level on liver and serum lipid metabolism of rats. Agricultural and Biological Chemistry. 46(3):703–707.

43. Keil-Stietz, K., & Lein, P. J. (2023). Gene×environment interactions in autism spectrum disorders. Current topics in developmental biology, 152, 221–284.

44. Keil Stietz KP, Sethi S, Klocke CR, de Ruyter TE, Wilson MD, Pessah IN, Lein PJ. 2021. Sex and genotype modulate the dendritic effects of developmental exposure to a human-relevant polychlorinated biphenyls mixture in the juvenile mouse. Front Neurosci. 15:766802.

45. Kelvington B, Kim J, Banerjee S, Stevens HE, Chow KH, Abel T. 2026. Disruption of DNA repair as an emerging epigenetic mechanism underlying autism spectrum disorder. Curr Psychiatry Rep. 28(1).

46. Kim MS, Kim Y, Choi H, Kim W, Park S, Lee D, Kim DK, Kim HJ, Choi H, Hyun DW et al. 2020. Transfer of a healthy microbiota reduces amyloid and tau pathology in an alzheimer’s disease animal model. Gut. 69(2):283–294.

47. Kim N, Jeon SH, Ju IG, Gee MS, Do J, Oh MS, Lee JK. 2021. Transplantation of gut microbiota derived from alzheimer’s disease mouse model impairs memory function and neurogenesis in c57bl/6 mice. Brain Behav Immun. 98:357–365.

48. Klaassen CD, Aleksunes LM. 2010. Xenobiotic, bile acid, and cholesterol transporters: Function and regulation. Pharmacol Rev. 62(1):1–96.

49. Klaassen CD, Cui JY. 2015. Review: Mechanisms of how the intestinal microbiota alters the effects of drugs and bile acids. Drug Metab Dispos. 43(10):1505–1521.

50. Klemm RW, Carvalho P. 2024. Lipid droplets big and small: Basic mechanisms that make them all. Annual Review of Cell and Developmental Biology. 40(Volume 40, 2024):143–168.

51. Koppel N, Maini Rekdal V, Balskus EP. 2017. Chemical transformation of xenobiotics by the human gut microbiota. Science. 356(6344).

52. Kostyniak PJ, Hansen LG, Widholm JJ, Fitzpatrick RD, Olson JR, Helferich JL, Kim KH, Sable HJ, Seegal RF, Pessah IN et al. 2005. Formulation and characterization of an experimental pcb mixture designed to mimic human exposure from contaminated fish. Toxicol Sci. 88(2):400–411.

53. Kwong E, Li Y, Hylemon PB, Zhou H. 2015. Bile acids and sphingosine-1-phosphate receptor 2 in hepatic lipid metabolism. Acta Pharm Sin B. 5(2):151–157.

54. Lein PJ, Yang D, Bachstetter AD, Tilson HA, Harry GJ, Mervis RF, Kodavanti PR. 2007. Ontogenetic alterations in molecular and structural correlates of dendritic growth after developmental exposure to polychlorinated biphenyls. Environ Health Perspect. 115(4):556–563.

55. Li N, Chen H, Cheng Y, Xu F, Ruan G, Ying S, Tang W, Chen L, Chen M, Lv L et al. 2021. Fecal microbiota transplantation relieves gastrointestinal and autism symptoms by improving the gut microbiota in an open-label study. Front Cell Infect Microbiol. 11:759435.

56. Li N, Wang Q, Wang Y, Sun A, Lin Y, Jin Y, Li X. 2019. Fecal microbiota transplantation from chronic unpredictable mild stress mice donors affects anxiety-like and depression-like behavior in recipient mice via the gut microbiota-inflammation-brain axis. Stress. 22(5):592–602.

57. Li Q, de Oliveira Formiga R, Puchois V, Creusot L, Ahmad AH, Amouyal S, Campos-Ribeiro MA, Zhao Y, Harris DMM, Lasserre F et al. 2025. Microbial metabolite indole-3-propionic acid drives mitochondrial respiration in cd4+ t cells to confer protection against intestinal inflammation. Nature Metabolism. 7(12):2510–2530.

58. Li X, Suh YP, Wilson RJ, Lein PJ, Cui JY, Lehmler HJ. 2024. Significant metabolic alterations in mouse dams exposed to an environmental mixture of polychlorinated biphenyls (pcbs) during gestation and lactation: Insights into pcb and metabolite profiles. Environ Toxicol Pharmacol. 111:104567.

59. Lim JJ, Goedken M, Jin Y, Gu H, Cui JY. 2024. Single-cell transcriptomics unveiled that early life bde-99 exposure reprogrammed the gut-liver axis to promote a proinflammatory metabolic signature in male mice at late adulthood. Toxicol Sci. 200(1):114–136.

60. Liu J, Tan Y, Song E, Song Y. 2020. A critical review of polychlorinated biphenyls metabolism, metabolites, and their correlation with oxidative stress. Chem Res Toxicol. 33(8):2022–2042.

61. Lyall K, Croen LA, Sjodin A, Yoshida CK, Zerbo O, Kharrazi M, Windham GC. 2017. Polychlorinated biphenyl and organochlorine pesticide concentrations in maternal mid-pregnancy serum samples: Association with autism spectrum disorder and intellectual disability. Environ Health Perspect. 125(3):474–480.

62. Megson, D., Idowu, I. G., & Sandau, C. D. (2024). Is current generation of polychlorinated biphenyls exceeding peak production of the 1970s?. The Science of the total environment, 924, 171436.

63. Mesnage R, Biserni M, Balu S, Frainay C, Poupin N, Jourdan F, Wozniak E, Xenakis T, Mein CA, Antoniou MN. 2018. Integrated transcriptomics and metabolomics reveal signatures of lipid metabolism dysregulation in heparg liver cells exposed to pcb 126. Arch Toxicol. 92(8):2533–2547.

64. Mesnier A, Champion S, Louis L, Sauzet C, May P, Portugal H, Benbrahim K, Abraldes J, Alessi M-C, Amiot-Carlin M-J et al. 2015. The transcriptional effects of pcb118 and pcb153 on the liver, adipose tissue, muscle and colon of mice: Highlighting of glut4 and lipin1 as main target genes for pcb induced metabolic disorders. PLoS One. 10(6):e0128847.

65. Monteiro-Cardoso VF, Corliano M, Singaraja RR. 2021. Bile acids: A communication channel in the gut-brain axis. Neuromolecular Med. 23(1):99–117.

66. Mouat, J. S., Li, X., Neier, K., Zhu, Y., Mordaunt, C. E., La Merrill, M. A., Lehmler, H. J., Jones, M. P., Lein, P. J., Schmidt, R. J., & LaSalle, J. M. (2023). Networks of placental DNA methylation correlate with maternal serum PCB concentrations and child neurodevelopment. Environmental research, 220, 115227.

67. Owe-Larsson M, Drobek D, Iwaniak P, Kloc R, Urbanska EM, Chwil M. 2025. Microbiota-derived tryptophan metabolite indole-3-propionic acid-emerging role in neuroprotection. Molecules. 30(17).

68. Park SH, Lee JH, Shin J, Kim JS, Cha B, Lee S, Kwon KS, Shin YW, Choi SH. 2021. Cognitive function improvement after fecal microbiota transplantation in alzheimer’s dementia patient: A case report. Curr Med Res Opin. 37(10):1739–1744.

69. Phillips JD. 2019. Heme biosynthesis and the porphyrias. Molecular Genetics and Metabolism. 128(3):164–177.

70. Powers BE, Widholm JJ, Lasky RE, Schantz SL. 2006. Auditory deficits in rats exposed to an environmental pcb mixture during development. Toxicol Sci. 89(2):415–422.

71. Premoli M, Pietropaolo S, Wöhr M, Simola N, Bonini SA. 2023. Mouse and rat ultrasonic vocalizations in neuroscience and neuropharmacology: State of the art and future applications. Eur J Neurosci. 57(12):2062–2096.

72. Pu Y, Zhang Q, Tang Z, Lu C, Wu L, Zhong Y, Chen Y, Hashimoto K, Luo Y, Liu Y. 2022. Fecal microbiota transplantation from patients with rheumatoid arthritis causes depression-like behaviors in mice through abnormal t cells activation. Transl Psychiatry. 12(1):223.

73. Ramstedt B, Slotte JP. 2002. Membrane properties of sphingomyelins. FEBS Lett. 531(1):33–37.

74. Ren X, Zou L, Zhang X, Branco V, Wang J, Carvalho C, Holmgren A, Lu J. 2017. Redox signaling mediated by thioredoxin and glutathione systems in the central nervous system. Antioxid Redox Signal. 27(13):989–1010.

75. Rieger MA, Dougherty JD. 2016. Analysis of within subjects variability in mouse ultrasonic vocalization: Pups exhibit inconsistent, state-like patterns of call production. Front Behav Neurosci. 10:182.

76. Rude, K. M., Pusceddu, M. M., Keogh, C. E., Sladek, J. A., Rabasa, G., Miller, E. N., Sethi, S., Keil, K. P., Pessah, I. N., Lein, P. J., & Gareau, M. G. (2019). Developmental exposure to polychlorinated biphenyls (PCBs) in the maternal diet causes host-microbe defects in weanling offspring mice. Environmental pollution (Barking, Essex : 1987), 253, 708–721.

77. Sable HJ, Powers BE, Wang VC, Widholm JJ, Schantz SL. 2006. Alterations in drh and drl performance in rats developmentally exposed to an environmental pcb mixture. Neurotoxicol Teratol. 28(5):548–556.

78. Saghazadeh A, Ataeinia B, Keynejad K, Abdolalizadeh A, Hirbod-Mobarakeh A, Rezaei N. 2019. Anti-inflammatory cytokines in autism spectrum disorders: A systematic review and meta-analysis. Cytokine. 123:154740.

79. Santini E, Klann E. 2014. Reciprocal signaling between translational control pathways and synaptic proteins in autism spectrum disorders. Sci Signal. 7(349):re10.

80. Seegal RF, Bush B, Brosch KO. 1994. Decreases in dopamine concentrations in adult, non-human primate brain persist following removal from polychlorinated biphenyls. Toxicology. 86(1-2):71–87.

81. Sethi S, Keil Stietz KP, Valenzuela AE, Klocke CR, Silverman JL, Puschner B, Pessah IN, Lein PJ. 2021. Developmental exposure to a human-relevant polychlorinated biphenyl mixture causes behavioral phenotypes that vary by sex and genotype in juvenile mice expressing human mutations that modulate neuronal calcium. Front Neurosci. 15:766826.

82. Sharon G, Cruz NJ, Kang DW, Gandal MJ, Wang B, Kim YM, Zink EM, Casey CP, Taylor BC, Lane CJ et al. 2019. Human gut microbiota from autism spectrum disorder promote behavioral symptoms in mice. Cell. 177(6):1600–1618 e1617.

83. Simola N, Granon S. 2019. Ultrasonic vocalizations as a tool in studying emotional states in rodent models of social behavior and brain disease. Neuropharmacology. 159:107420.

84. Sinioja T, Bodin J, Duberg D, Dirven H, Berntsen HF, Zimmer K, Nygaard UC, Orešič M, Hyötyläinen T. 2022. Exposure to persistent organic pollutants alters the serum metabolome in non-obese diabetic mice. Metabolomics. 18(11):87.

85. Sprowles JLN, Monaikul S, Aguiar A, Gardiner J, Monaikul N, Kostyniak P, Schantz SL. 2022. Associations of concurrent pcb and pbde serum concentrations with executive functioning in adolescents. Neurotoxicol Teratol. 92:107092.

86. Sredlova K, Cajthaml T. 2022. Recent advances in pcb removal from historically contaminated environmental matrices. Chemosphere. 287(Pt 1):132096.

87. Stewart PW, Reihman J, Lonky EI, Darvill TJ, Pagano J. 2003. Cognitive development in preschool children prenatally exposed to pcbs and mehg. Neurotoxicol Teratol. 25(1):11–22.

88. Studer E, Zhou X, Zhao R, Wang Y, Takabe K, Nagahashi M, Pandak WM, Dent P, Spiegel S, Shi R et al. 2012. Conjugated bile acids activate the sphingosine-1-phosphate receptor 2 in primary rodent hepatocytes. Hepatology. 55(1):267–276.

89. Sumra B, Kocherry C, Shamim H, Jhakri K, Al-Shudifat M, Mohammed L. 2025. Impact of omega-3 fatty acids on cognitive outcomes in children with autism spectrum disorder: A systematic review. Cureus. 17(3):e80291.

90. Valente M, Dentoni M, Bellizzi F, Kuris F, Gigli GL. 2022. Specialized pro-resolving mediators in neuroinflammation: Overview of studies and perspectives of clinical applications. Molecules. 27(15).

91. Vogel C, Marcotte EM. 2012. Insights into the regulation of protein abundance from proteomic and transcriptomic analyses. Nat Rev Genet. 13(4):227–232.

92. Wahlang B, Hardesty JE, Jin J, Falkner KC, Cave MC. 2019. Polychlorinated biphenyls and nonalcoholic fatty liver disease. Current Opinion in Toxicology. 14:21–28.

93. Wan L, Wang H, Liang Y, Zhang X, Yao X, Zhu G, Cai J, Liu G, Liu X, Niu Q et al. 2024. Effect of oral faecal microbiota transplantation intervention for children with autism spectrum disorder: A randomised, double-blind, placebo-controlled trial. Clin Transl Med. 14(9):e70006.

94. Wang T, Chen B, Luo M, Xie L, Lu M, Lu X, Zhang S, Wei L, Zhou X, Yao B et al. 2023. Microbiota-indole 3-propionic acid-brain axis mediates abnormal synaptic pruning of hippocampal microglia and susceptibility to asd in iugr offspring. Microbiome. 11(1):245.

95. Wayman GA, Yang D, Bose DD, Lesiak A, Ledoux V, Bruun D, Pessah IN, Lein PJ. 2012. Pcb-95 promotes dendritic growth via ryanodine receptor-dependent mechanisms. Environ Health Perspect. 120(7):997–1002.

96. Weber R, Herold C, Hollert H, Kamphues J, Ungemach L, Blepp M, Ballschmiter K. 2018. Life cycle of pcbs and contamination of the environment and of food products from animal origin. Environ Sci Pollut Res Int. 25(17):16325–16343.

97. Wen TH, Binder DK, Ethell IM, Razak KA. 2018. The perineuronal ’safety’ net? Perineuronal net abnormalities in neurological disorders. Front Mol Neurosci. 11:270.

98. Wilson RJ, Suh YP, Dursun I, Li X, da Costa Souza F, Grodzki AC, Cui JY, Lehmler HJ, Lein PJ. 2024. Developmental exposure to the fox river pcb mixture modulates behavior in juvenile mice. Neurotoxicology. 103:146–161.

99. Yang D, Kim KH, Phimister A, Bachstetter AD, Ward TR, Stackman RW, Mervis RF, Wisniewski AB, Klein SL, Kodavanti PR et al. 2009. Developmental exposure to polychlorinated biphenyls interferes with experience-dependent dendritic plasticity and ryanodine receptor expression in weanling rats. Environ Health Perspect. 117(3):426–435.

100. Yang D, Lein PJ. 2010. Polychlorinated biphenyls increase apoptosis in the developing rat brain. Curr Neurobiol. 1(1):70–76.

101. Yin X, Chen L, Xia Y, Cheng Q, Yuan J, Yang Y, Wang Z, Wang H, Dong J, Ding Y et al. 2016. Maternal deprivation influences pup ultrasonic vocalizations of c57bl/6j mice. PLoS One. 11(8):e0160409.

102. Zeidan J, Fombonne E, Scorah J, Ibrahim A, Durkin MS, Saxena S, Yusuf A, Shih A, Elsabbagh M. 2022. Global prevalence of autism: A systematic review update. Autism Res. 15(5):778–790.

