## Supplemental Materials for "PCB-Exposed Gut Microbiota Recapitulate Neurobehavioral Deficits and Reprogram Hippocampal Transcriptomic Signatures"

**Untargeted Metabolomics Sample Preparation and LC-MS/MS Analysis**

Serum samples were thawed on ice and vortexed for 10 seconds. A 50 μL aliquot of each sample was extracted with 300 μL of acetonitrile:methanol (1:4, v/v) containing internal standards in a 2 mL microcentrifuge tube. Samples were vortexed for 3 min and centrifuged at 12,000 rpm for 10 min at 4 °C. An aliquot (200 μL) of the supernatant was collected and incubated at -20 °C for 30 min to enhance precipitation, followed by an additional centrifugation step at 12,000 rpm for 3 min at 4 °C. A final volume of 180 μL supernatant was collected for LC-MS analysis. Chromatographic separation was performed using a Waters ACQUITY Premier HSS T3 column (1.8 μm, 2.1 × 100 mm) on an ExionLC AD system (SCIEX) with a mobile phase consisting of ultrapure water with 0.1% formic acid (phase A) and acetonitrile with 0.1% formic acid (phase B). The column temperature was maintained at 40 °C, with a flow rate of 0.40 mL/min and an injection volume of 2 μL, delivered over a 14-minute gradient. Mass spectrometric detection was performed on a TripleTOF 6600+ instrument (SCIEX) equipped with an electrospray ionization source operated in both positive and negative ion modes, with data acquired in information-dependent acquisition mode across an MS1 range of 50–1000 Da and an MS2 range of 25–1000 Da.

**Untargeted Metabolomics Data Processing and Quality Control**

Raw LC-MS data were converted to mzML format using ProteoWizard, and peak detection, alignment, and retention time correction were performed using XCMS. Features with a missing rate greater than 50% within any experimental group were excluded. Missing values were imputed using a combination of k-nearest neighbors (KNN) and minimum value substitution (1/5 minimum), and signal drift was corrected using support vector regression (SVR). Metabolite annotation was achieved through comparison with in-house and public databases, as well as predictive tools including metDNA. Only metabolites with a comprehensive identification score >0.5 and a coefficient of variation (CV) <0.3 in quality control (QC) samples were retained. Data from positive and negative ionization modes were merged, prioritizing features with the highest identification confidence and lowest variability, yielding a total of 6,472 identified metabolites, of which 4,125 were assigned MS2-based structural annotations. Quality control samples were generated by pooling aliquots from all study samples and were injected at regular intervals (every 10 samples) throughout data acquisition. Instrument stability and reproducibility were assessed by evaluating the overlap of total ion chromatograms (TICs) across QC injections and by calculating Pearson correlation coefficients between QC samples. Data quality was further evaluated using CV distribution analysis, with high proportions of metabolites exhibiting CV values <0.5 and <0.3 indicating stable and highly reproducible analytical performance.

**Supplemental Figure 1. Effects of PCB-associated FMT on neonatal ultrasonic vocalizations, body weight, and body temperature in P7 offspring. (A)** Litter-averaged ultrasonic vocalizations (USVs) recorded during 3 minutes of maternal separation in male and female P7 offspring following PCB-associated FMT compared with vehicle-associated FMT. Each data point represents the average USV count within a litter, with bars showing group means and error bars. **(B)** Body weight distributions of male and female P7 offspring in vehicle-FMT and PCB-associated FMT groups. Violin plots display individual measurements and their distributions. Significant differences between indicated groups are shown (*P* < 0.0001 and *P* = 0.0007). **(C)** Litter-averaged body temperature in male and female P7 offspring following vehicle-associated or PCB-associated FMT. Violin plots show the distributions of litter-level measurements. **(D-E)** Simple linear regression analyses examining the associations between USV counts and body weight (D; R² = 0.0398, *P* = 0.2904) or body temperature (E; R² = 0.0007, *P* = 0.8888). Solid lines indicate fitted linear regression relationships. Sex- and FMT-associated effects on USVs were evaluated using two-way analysis of variance (ANOVA), with litter averages serving as the biological units for USV analyses. Simple linear regression was used to assess associations between USV counts and body weight or temperature. Reported *P* values are indicated in the
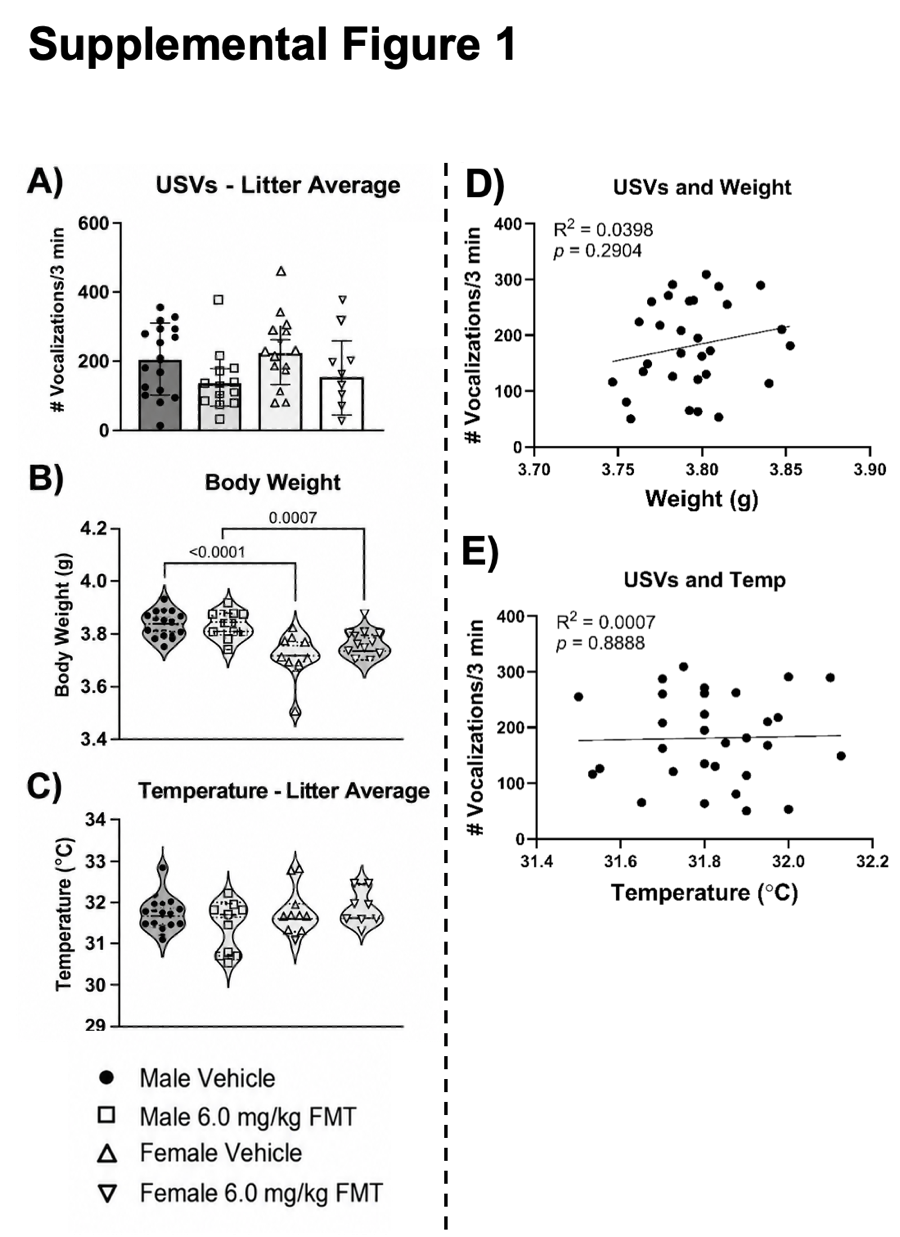
corresponding panels.
